# A comprehensive comparison on cell type composition inference for spatial transcriptomics data

**DOI:** 10.1101/2022.02.20.481171

**Authors:** Jiawen Chen, Weifang Liu, Tianyou Luo, Zhentao Yu, Minzhi Jiang, Jia Wen, Gaorav P. Gupta, Paola Giusti, Hongtu Zhu, Yuchen Yang, Yun Li

**Author notes:** These authors contributed equally (Jiawen Chen, Weifang Liu, Tianyou Luo). Corresponding author: Yun Li, Department of Genetics, 120 Mason Farm Road, Campus Box 7264, University North Carolina, Chapel Hill, NC 27599, USA.

## Abstract

Spatial transcriptomic (ST) technologies allow researchers to examine high-quality RNA-sequencing data along with maintained two-dimensional positional information as well as a co-registered histology image. A popular use of ST omics data is to provide insights about tissue structure and spatially unique features. However, due to the technical nature unique to most ST data, the resolution varies from a diameter of 2-10*μm* to 50-100*μm* instead of single-cell resolution, which brings uncertainty into cell number and cell mixture within each ST spot. Motivated by the important role for spatial arrangement of cell types within a tissue in physiology and disease pathogenesis, several ST deconvolution methods have been developed and are being used to explore gene expression variation and identification of spatial domains. The aim of this work is to review state-of-the-art methods for ST deconvolution, while comparing their strengths and weaknesses. Specifically, we use four real datasets to examine the performance of eight methods across different tissues and technological platforms.

**Key Points:**

- Cell mixture inference is a critical step in the analysis of spatial transcriptomics (ST) data to prevent downstream analysis suffering from confounding factors at the spot level.
- Existing ST deconvolution methods can be classified into three groups: probabilistic-based, non-negative matrix factorization and non-negative least squares based, and other deep learning framework-based methods.
- We compared eight ST deconvolution methods by using two single cell level resolution datasets and two spot level resolution ST datasets. We provided practical guidelines for the choice of method under different scenarios as well as the optimal subsets of genes to use for each method.

## Introduction

Analysis of patterns of messenger RNAs in a histological tissue section is a cornerstone of biomedical research and diagnostics. In recent years, we have witnessed rapid development and advances of novel spatial transcriptomic (ST) technology [1–4]. By labeling histological sections with unique positional barcodes, most ST technology can obtain high-quality RNA-sequencing data, while maintaining two-dimensional positional information as well as a co-registered histology image [1–4]. ST technology can be classified into two categories, including imaging- and sequencing-based methods [5]. Imaging-based techniques, including single-molecule fluorescence in situ hybridization (smFISH) [6] and multiplexed error robust fluorescence in situ hybridization (MERFISH) [1], provide both quantitative measurements of RNA expression levels and information about RNA spatial localization by directly imaging individual RNA molecules in singlecells. Sequencing-based techniques, such as spatial barcoding employed by the widely-used 10X Genomics Visium platform, is powered by spatially barcoded mRNA-binding oligonucleotides to capture gene expression in tissue samples [4].

The invention of ST technology offers a more powerful and efficient way to explore the existence of spatial transcriptomics pattern, including variance in gene expression, cellular subpopulation, and colocalization with a much higher resolution than prior methods. Such spatial patterns compose the organ system and deeply intertwine with organ structure and function. Technologies like single-cell RNA sequencing (scRNA-seq) often require isolation by using either fluorescence-activated cell sorting (FACS) or manual picking to study organ structure, such as the layer-specific enrichment patterns [7, 8]. Such isolation can be labor-intensive and costly. Instead, ST technologies are able to reveal such diversity of molecularly defined cell types in a single experiment [2].

However, existing ST techniques are limited in the trade-off between spatial resolution and the number of genes measured. Imaging-based ST techniques can obtain single cell or even subcellular level data, but most of them can only measure around 100-1000 genes, entailing *a priori* marker gene selection and rendering them less suitable for exploratory analyses. Sequencing-based techniques can measure whole-transcriptome-wise gene expression, whereas these technologies can only obtain spot-level data (diameter 2-10 or 50-100 *μm*) that approaches, but does not achieve, single-cell resolution. Therefore, any downstream analysis could potentially suffer from confounding factors at the spot level. Specifically, gene expression variation across spots could be driven by features like the different mixture of cell types and the number of cells as opposed to actually being driven by spatial location. To appropriately adjust for confounding factors at the spot level, ST deconvolution methods have been recently developed for more confidently revealing underlying biological mechanisms. To make sound scientific inferences, we need a fair assessment of various state-of-the-art ST deconvolution methods. In the literature [9–18], the performance of each deconvolution method has been assessed predominantly by the developers, using simulated datasets with varied assumptions presented in different studies. These comparisons could be incomplete and prone to biases in interpretation. There are two types of simulated datasets. The first kind of simulation is to select cells from real scRNA-seq datasets and combine their transcriptomic profiles to mimic ST spots[18]. In some other studies, such comparisons are performed on simulated scRNA-seq and ST datasets, where the gene expression is simulated from Poisson, Gaussian, negative binomial, Gamma distribution, etc. with certain additional assumptions, such as co-localization of several cell types or the existence of spatial varying gene expression patterns[10]. Both types of simulated datasets have disadvantages as they fail to reflect a wider spectrum of realistic data, are unrealistic at least in some major aspects, and favor the “promoted” method developed by the authors of the corresponding publication. To the best of our knowledge, no third-party comprehensive evaluation of the ST deconvolution methods has been performed using the same set of diverse real datasets. In this work, we aim to fill in this gap.

In this review and analysis, we summarize and compare computational strategies commonly employed for cell type deconvolution for ST data. The review is organized as follows. We first describe the commonly used ST deconvolution methods, highlighting several key aspects including the employed statistical method, type(s) of ST data tailored to, and method-specific unique features. We then present performance of these methods using the same four real ST datasets as benchmarks. Finally, we provide practical guidelines and emphasize the advantages and drawbacks of these methods in real data applications.

### Computational methods developed for cell type deconvolution of ST data

In recent years, a diverse collection of ST deconvolution algorithms has been proposed. Existing deconvolution methods for ST data can be largely classified into three groups: probabilistic methods, methods based on non-negative matrix factorization (NMF) and non-negative least squares (NNLS), and deep learning-based methods (**Figure 1, Table 1**). The first group of probabilistic methods includes Adroit [16], Cell2location [18], RCTD [9], STdeconvolve [17], and Stereoscope [11], where the data distribution is explicitly or parametrically specified and inference is carried out based on model likelihood. The second group of NMF and NNLS based methods includes SPOTlight [15] and spatialDWLS [14]. The third group of deep learning-based methods, such as, DSTG [12] and Tangram [13], optimizes a specifically designed loss function (**Figure 1, Table 1**). Here we review eight state-of-the-art methods: Adroit, Cell2location, DestVI, RCTD, STdeconvolve, Stereoscope, SPOTlight, and DSTG [9–12, 15–18]. We briefly summarize each method below.

**Figure 1.**
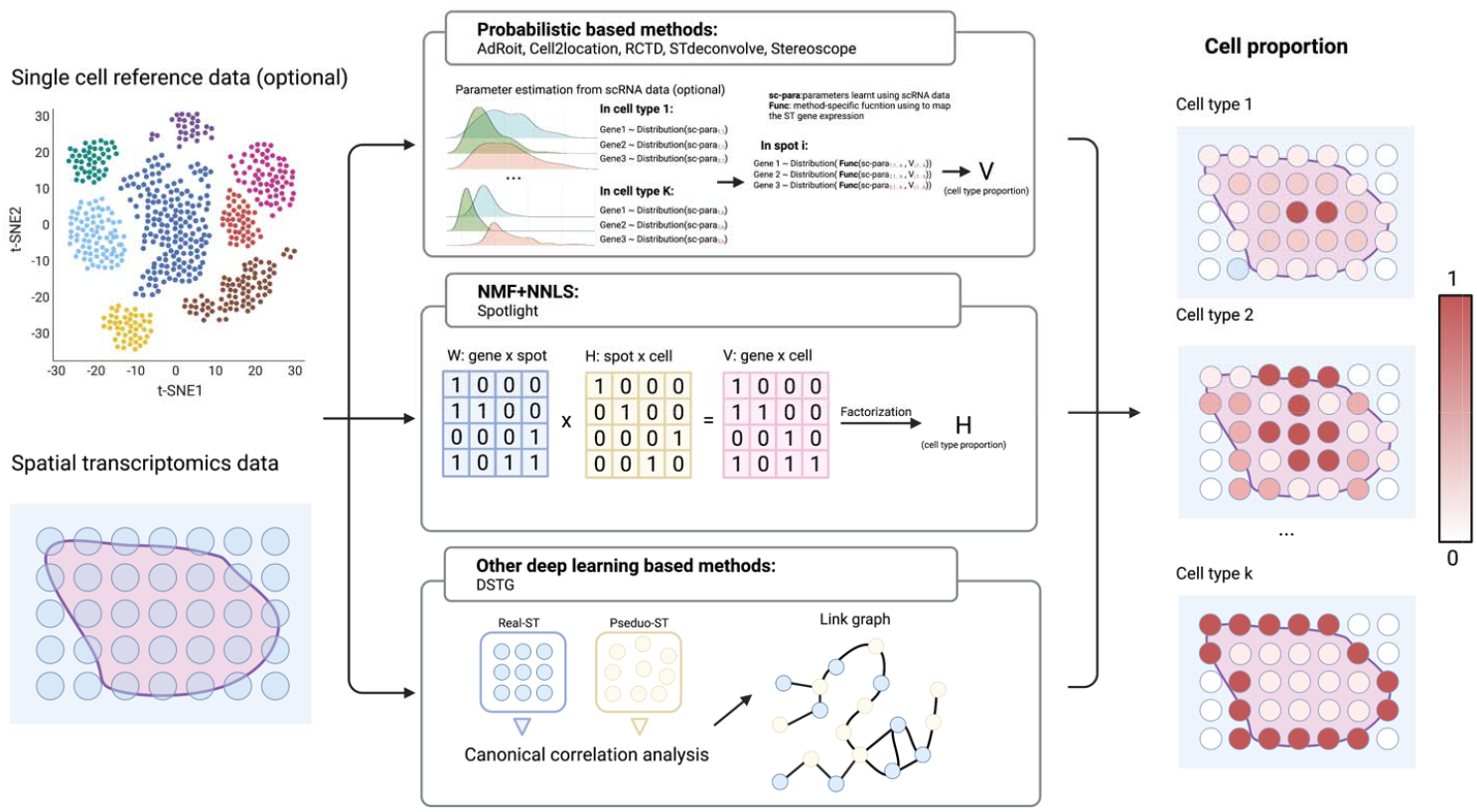
Summary of ST deconvolution methods. ST deconvolution methods take ST data and an optional scRNA-seq reference data as input (left panel). Current ST deconvolution methods can be classified into three main categories: probabilistic-based, NMF and NNLS based, and deep learning methods (center panel). The output of ST deconvolution methods is cell proportion for each spot which scales from 0 to 1 (right panel).

**Table 1.** ST deconvolution methods overview.

| Method | Designed for ST data? | Feature selection | Inference method | Language | URLs | Reference |
| --- | --- | --- | --- | --- | --- | --- |
| AdRoit | No | Genes enriched in one or more cell types or highly variable genes | Probabilistic, non-negative least squares regression | R | https://github.com/TaoYang-dev/AdRoit | [16] |
| Cell2location | Yes | No selection | Probabilistic, negative binomial distribution, variational Bayesian inference | Python | https://cell2location.readthedocs.io/en/latest/ | [18] |
| DSTG | Yes | 2,000 most variable genes | Semi-supervised graph convolutional network, adaptive moment estimation algorithm | Python | https://github.com/Su-informatics-lab/DSTG | [12] |
| DestVI | Yes | Highly variable genes | Probabilistic, latent variable models, auto-encoding variational bayes | Python | https://docs.scvi-tools.org/en/stable/user_guide/models/destvi.html | [10] |
| RCTD | Yes | DE genes | Probabilistic, Poisson distribution, maximum likelihood | R | https://github.com/dmcable/spacexr | [9] |
| SPOTlight | Yes | Highly variable genes | Non-negative matrix factorization (NMF) along with non-negative least squares (NNLS) | R | https://github.com/MarcElosua/SPOTlight_deconvolution_analysis | [15] |
| STdeconvolve | Yes | Highly variable genes | Generative probabilistic model: latent Dirichlet allocation (LDA), variational expectation-maximization algorithm | R | https://jef.works/STdeconvolve/ | [17] |
| Stereoscope | Yes | Top 5000 highest expressed genes (optional) | Probabilistic, negative binomial distribution | Python | https://github.com/almaan/Stereoscope | [11] |

Accurate and Robust Method to Infer Transcriptome Composition (AdRoit) [16] is designed for bulk RNA-seq data, but it can also be used for spatially resolved transcriptomics data. AdRoit first selects informative genes and then estimates their corresponding locations and dispersion parameters of negative binomial distribution by assuming that counts follow negative binomial distributions. Subsequently, cross-sample variability, collinearity of expression profiles, and cell type specificity of cell types are estimated from multi-cell resolution ST data. Then, the gene-wise scaling factors are estimated using both scRNA-seq data and ST data. Finally, these quantities are included in a weighted regularized model for inferring cell-type proportions. The key development of AdRoit is that it adopts a learning approach to correct sequencing platform bias on genes by using gene-wise scaling factors.

Cell2location [18] adopts a hierarchical Bayesian framework that assumes gene expression counts follow a negative binomial distribution. It first uses external scRNA-seq data as reference to estimate cell type specific signatures. The observed spatial expression count matrix is modeled with a negative binomial distribution, where gene-specific technological sensitivity, gene- and location-specific additive shifts are included as part of the mean parameter. Then cell2location uses variational Bayesian inference to approximate the posterior distribution and produces parameter estimates accordingly.

Deconvolution of Spatial Transcriptomics profiles using Variational Inference (DestVI) [10] is a probabilistic method for multi-resolution analysis of ST data. DestVI explicitly models variation within cell types via continuous latent variables instead of limiting the analysis to a discrete view of cell types. Such continuous within-cell-type variations as well as the corresponding cell-type-specific profiles are learned through a conditional deep generative model, specifically, using variational inference with decoder neural networks. In this scheme, two different latent variable models (LVMs) are constructed for scRNA-seq (scLVM) and ST data (stLVM), respectively. DestVI similarly assumes that the number of observed transcripts follows a negative binomial distribution. The decoder neural network trained by scLVM is employed by stLVM and cell type proportion is obtained using maximum-a-posteriori (MAP) inference scheme where the number of observed transcripts in each spot is assumed to follow a weighted sum of the inferred single-cell negative binomial distributions.

RCTD [9] is initially designed for Slide-seq data, but it could also be used on other ST data. It assumes that the observed spot-level gene counts for each spot follow a Poisson distribution, while accounting for platform effects by including a gene-specific random effect term. RCTD first uses external scRNA-seq reference data to estimate the mean gene expression profile of each cell type. Gene filtering is then performed by selecting differentially expressed (DE) genes across cell types, and the variance of gene-specific platform effects is estimated. The inferred platform effects are plugged into the probabilistic model to obtain the maximum likelihood estimates (MLE) of cell type proportions.

STdeconvolve [17] is a reference-free and unsupervised cell type deconvolution method for ST data. The key difference between STdeconvolve and other methods is that STdeconvolve can perform cell type deconvolution without using external scRNA-seq references. The method is built upon latent Dirichlet allocation (LDA) to identify putative transcriptional profiles for each cell type and their proportions in each ST spot, which is defined as a mixture of a predetermined number of cell types modeled by a multinomial distribution and cell type distribution is drawn from a uniform Dirichlet distribution. STdeconvolve assumes the existence of highly co-expressed genes for each cell type and selects significantly over-dispersed genes to inform latent cell types.

Stereoscope [11] achieves the deconvolution purpose, while spatially mapping cell types using annotated scRNA-seq reference and ST data. Stereoscope also relies on the commonly used assumption that counts from both spatial and single-cell data follow a negative binomial distribution. Stereoscope includes a noise term as a form of “dummy” cell type to account for asymmetric datasets where the cell types do not overlap perfectly in spatial and single-cell data. MLE is employed to estimate cell type specific parameters using scRNA-seq reference data and MAP is used to infer cell type mixture in the ST data.

SPOTlight [15] is a deconvolution algorithm that employs the non-negative matrix factorization (NMF) regression algorithm as well as the non-negative least squares (NNLS). In SPOTlight, NMF is carried out to identify cell type-specific topic profiles in scRNA-seq references and NNLS is carried out to identify spot topic profiles, which is the deconvolution result. Besides, SPOTlight is reported to perform sensitively and accurately across different biological scenarios and varying technology versions with matched and external references.

DSTG [12] is a similarity-based semi-supervised graph convolutional network (GCN) model that can recover cell type proportions in each spot. By leveraging scRNA-seq data, DSTG first constructs a synthetic ST data called “pseudo-ST” by randomly pooling two to eight cells selected from scRNA-seq data as a ST spot. Then, to capture the similarity between spots and incorporate pseudo-ST and real ST data, DSTG learns a link graph by finding mutual nearest neighbors in the shared space identified by canonical correlation analysis (CCA). A semi-supervised GCN is trained with both the pseudo and real ST data and the link graph, which can be used to predict cell type proportions in real ST data.

In summary, many powerful ST deconvolution tools have been developed and tailored specifically for ST datasets. These methods have demonstrated their utility in both simulated and real datasets. By conducting downstream analysis with the knowledge of the cell proportion variation in ST data along with the spatial information, we can better uncover the underlying biological mechanism and further discover novel findings that we could not achieve using scRNA-seq datasets. However, there does not exist a fair and comprehensive comparison of these cell deconvolution methods yet. We will use multiple real ST datasets, encompassing both single-cell level resolution and spot-level resolution ST data with pathologist annotation (**Table 2**), to systematically and objectively evaluate the performance of these methods.

**Table 2.**
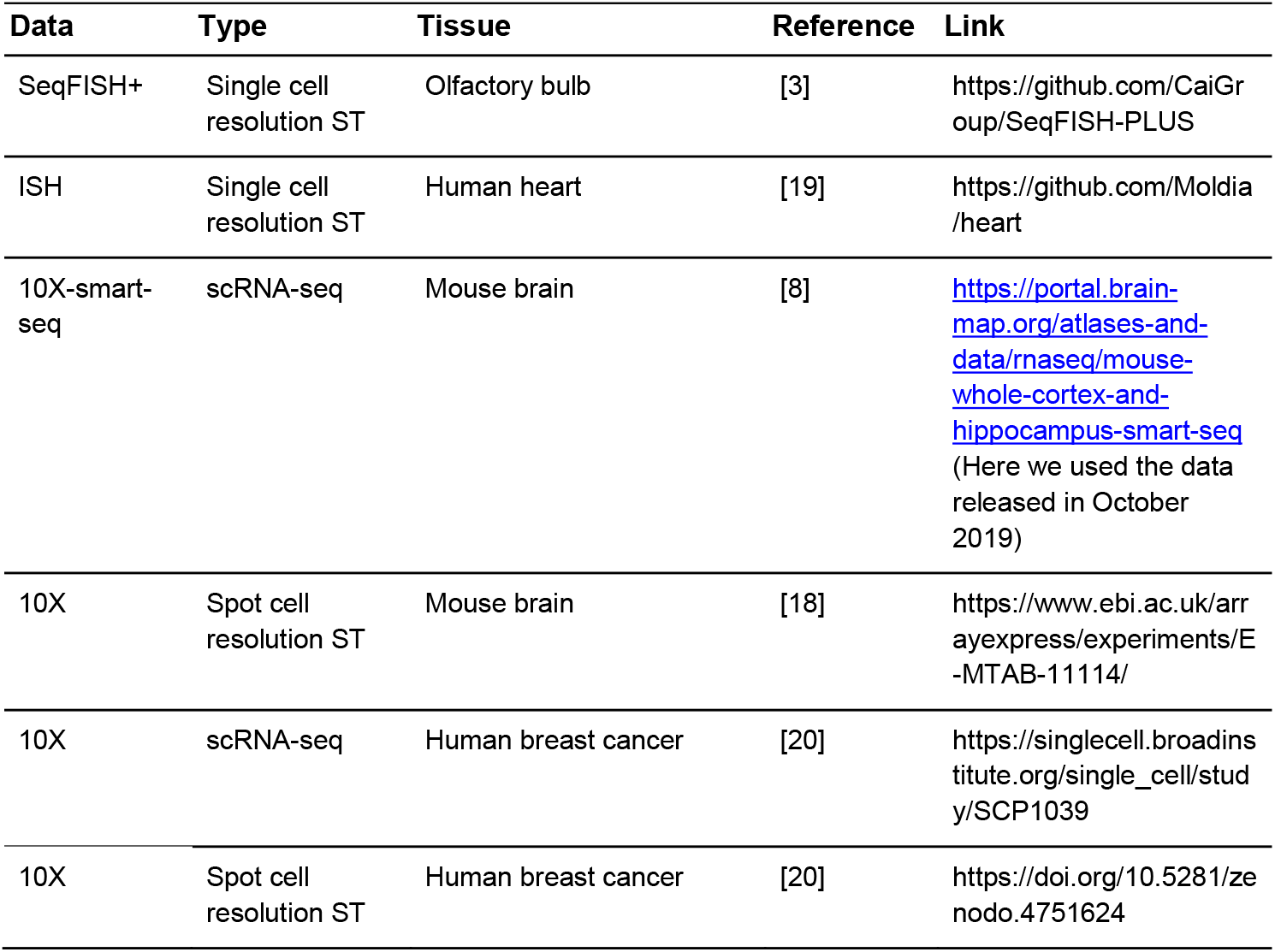
Data source and reference

### Benchmarking ST deconvolution methods performance

We employed four real datasets [3, 18–20] to evaluate the performance of the aforementioned eight methods [9–12, 15–18] (**Table 2**). We first used two ST datasets at single-cell level resolution [3, 19], which come from mouse olfactory bulb and human developing heart. We pooled the cells together according to their spatial coordinates to mimic ST spots (**Methods**). Then we assessed the cell type proportion estimates for each deconvolution method using RMSE (root mean square error) and average Pearson correlation across cell types. As RMSE is a distance metric, values closer to 0 signify a higher similarity between two distributions. On the other hand, a higher Pearson correlation indicates a better ability to identify cell type distribution patterns across spots. The other two ST datasets come from mouse brain [18] and human breast cancer [20] which have spot-level resolution. We first inferred cell type proportions for each spot and then clustered the spots using the estimated cell type proportions. Finally, we compared the clustering results with pathological annotation. We evaluated the performance of the deconvolution methods using the clustering results and quantified them using adjusted mutual information (AMI) (**Methods**).

### Evaluation with single-cell resolution ST data

We first evaluated the deconvolution methods on a SeqFISH+ dataset from mouse olfactory bulb tissue [3]. SeqFISH+ is capable of imaging mRNAs for 10,000 genes in single cells with high accuracy and sub-diffraction-limit resolution using a standard confocal microscope [3]. In this SeqFISH+ dataset, 7 fields of views (FOV) of olfactory bulb data were available and each FOV was cropped into 25 spots (**Figure 2A**). Only spots with non-zero cells were kept for analysis. For deconvolution methods that require a scRNA-seq reference, the olfactory bulb data itself was employed as the reference. The perfectly matching reference eliminates the potential performance impairment caused by any systematic differences between single-cell reference and ST data. Since the selection of genes in the analysis is critical to deconvolution performance, we used several gene subsets. A “default” gene set was subsetted using the built-in gene selection strategy of each method and therefore is method-specific (**Methods**). For methods without a specific recommendation or built-in gene selection, we used 2000 highly variable genes (HVG). We also selected the top 100, 500, and 1000 HVG using the single-cell reference (**Methods**) and used the overlapping genes in the ST data. Moreover, we subsetted another gene set containing the top 12, 70, 148 marker genes for each cell type (**Methods**) to make the number of unique genes summing approximately to 100, 500, and 1000 genes in total **(Figure 2B)**.

**Figure 2.**
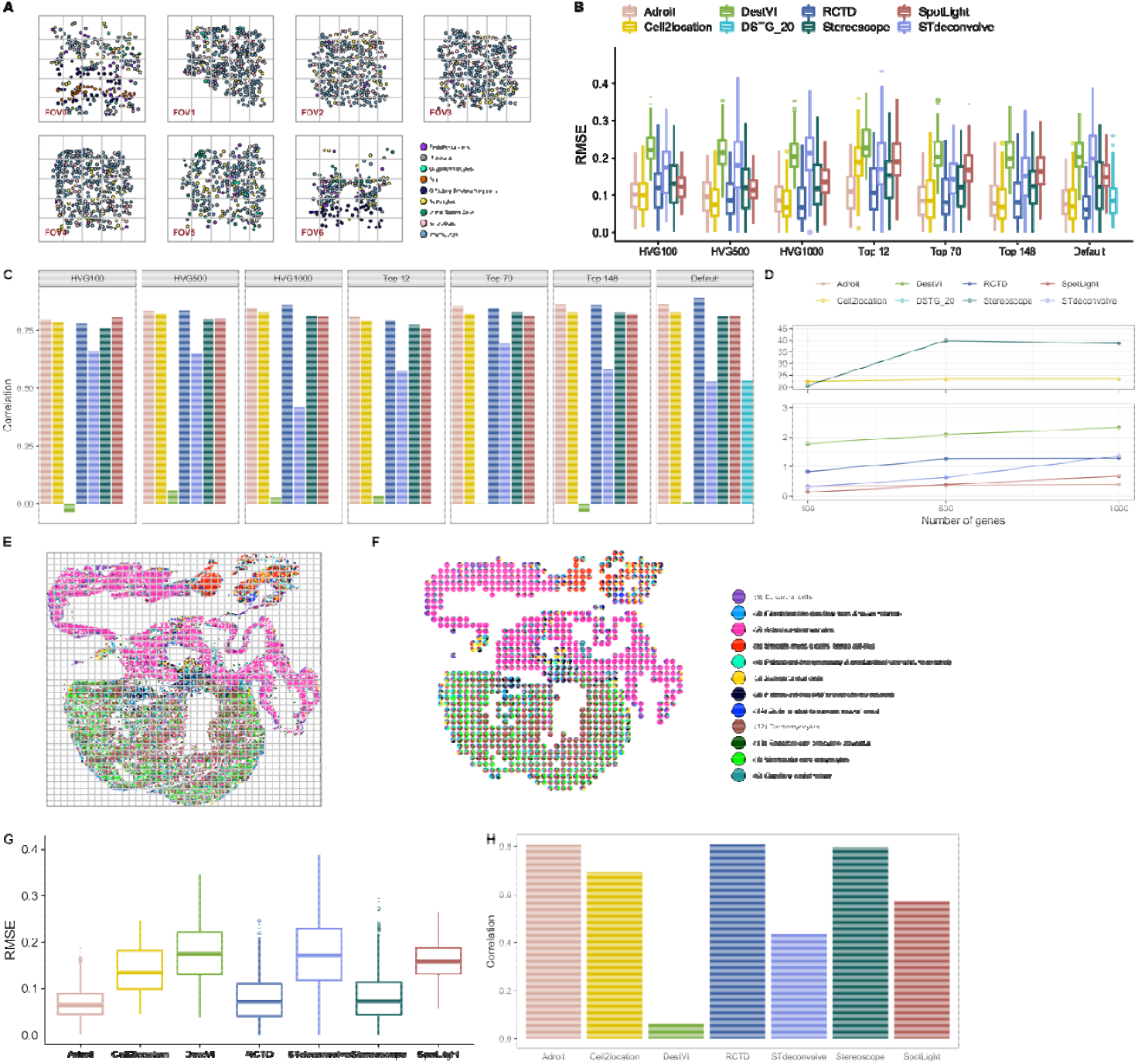
Evaluation on single-cell resolution ST data. (**A**) Overview of the cell atlas of 7 fields of view in the mouse olfactory bulb (OB) data set. (**B**) RMSE of OB cell proportion estimates from 8 methods using sets of highly variable genes (HVG), sets of marker genes (Top 12, Top 40, Top 148) and the method-specific default gene set (Default). (**C**) Pearson correlation of OB cell deconvolution results from 8 methods using the same sets of genes as in (B). (**D**) Computing time (in minutes) when using 100, 500, 1000 HVG including both single-cell inference (if present) and ST spot deconvolution. (**E**) Overview of the cell atlas in the developing human heart PCW6.5_1 data set. (**F**) Cell type proportion in each pooled spot. Each spot is a pie chart. (**G**) RMSE of heart cell proportion estimates from 8 methods using the methodspecific default gene set for the human heart dataset. (**H**) Pearson correlation of heart cell proportion estimates from 8 methods using the same sets of genes as in (G). Colors of the boxplots and bar plots follow the same labels and legend in (B) and (D).

We observed considerable low RMSE in Adroit, Cell2Location, RCTD, and Stereoscope inferred results (**Figure 2B**). RMSE consistently decreases as the number of genes increases from 100 to 1000 in both HVG and marker gene subsets for most methods. Employing the same number of genes, Cell2Location and SPOTlight perform better using HVG, whereas other methods exhibit mixed patterns among varying gene subsets. DSTG shows consistent results with different hyperparameters (k=20,50, and 100) with its default gene set (**Supplementary Figure 1**). As a side note, DSTG does not support inference for datasets with fewer than 1000 genes. The patterns observed above with the RMSE metric remain qualitatively similar when assessed with the Pearson correlation metric, which we calculated as the average correlation over all cell types (**Figure 2C**). Specifically, Adroit, Cell2Location, RCTD and Stereoscope inferred results also achieve high correlation regardless of the gene subsets. The correctly inferred proportion of Mitral/Tufted Cells is one of the reasons for the superior performance of these methods in FOV6 and FOV7. In the tSNE embedding space, Mitral/Tufted Cells have similar characteristics to Interneuron and Neuroblast. These similarities may explain why some methods miss the pattern (**Supplementary Figure 2**). SPOTlight also attains high correlation; however, the cell proportion estimates are much noisier than the four methods highlighted above (**Supplementary Figure 1**). STdeconvolve, as a reference-free method, entails cell type mapping after unsupervised inference. Instead of letting STdeconvolve infer the number of cell types, we used the true cell type numbers in the evaluation. In this case, STdeconvolve may not be able to achieve its best performance since the inaccurate cell type mapping may contribute to the poor performance of STdeconvolve (**Supplementary Figure 3, Methods**). Using transcriptional correlation, one STdeconvolve inferred cell type could have high correlation with multiple ground truth cell types. When this happens, if we decide to assign one ground truth cell type to one inferred cell type, then it is highly likely that some cell type(s) will be mismatched in terms of low correlation. For example, when using the default gene set, the inferred cell type 3 is labeled as Pia despite their correlation being −0.10 only because other ground truth cell types have higher correlation with other inferred cell types (**Supplementary Figure 3**).

We further benchmarked the runtime using the set of 100, 500, and 1000 highly variable genes (**Figure 2D**). The runtime increases linearly with the number of genes in the dataset for most of the methods. Adroit and SPOTlight are among the fastest methods with runtime less than a minute. The runtime of Cell2Location, Stereoscope and DestVI is heavily dependent on the number of training epochs (**Methods**). It is difficult to choose the optimal number of training epochs to prevent underfitting. A conservative solution is to increase the training epoch number, which will consequently increase the runtime. Examining the results for Stereoscope and Cell2location, we observe that their RMSE does not vary significantly with top 500 HVG, top 1000 HVG and the default gene set which is top 2000 HVG. The runtime is around 25 and 40 minutes when using top 1000 HVG for Stereoscope and Cell2location respectively. The runtime of DSTG with default gene set is around 2 minutes.

To evaluate the performance of the deconvolution methods with an extremely limited number of genes, we analyzed a dataset with merely 69 genes from the developing human heart (**Figure 2E**)[19]. This dataset is a subcellular resolution ST data developed by in situ sequencing (ISS). The spatial cell map and assignment of gene reads to individual cells were accomplished using ISS and pciSeq [21, 22]. According to the literature [19], the 69 genes consist of spatial marker genes identified in the previous ST analysis and marker genes for each major cluster from scRNA-seq analysis as well as genes previously reported to be important for cardiac development. Due to the limited number of genes, no further gene filtering was applied to methods without a gene filtering strategy besides the default data preprocessing procedure (**Methods**). The cells in the heart tissue were pooled into spots each of size 454*424 square pixel according to the spatial coordinates (**Figure 2F**).

Adroit, RCTD and Stereoscope retain their superior performance with the limited number of genes (**Figure 2G-H, Supplementary Figure 4**). They successfully map the atrial cardiomyocytes and ventricular cardiomyocytes into the atria and the ventricular body, respectively. Smooth muscle cells are also perfectly mapped to the outflow tract and epicardial cells are mapped to the thin outer layer of the heart, which all agree with annotations from the Cell Atlas (**Figure 2E, Supplementary Figure 4**) and previous studies [19, 23, 24]. Ventricular cardiomyocytes and cardiomyocytes exhibit similarity within the tSNE embedding space, as well as a co-localization pattern in the ventricular interval and the ventricle (**Figure 2E-F, Supplementary Figure 2**). Adroit fails to clearly capture the enriched pattern of cardiomyocytes in the ventricular interval. DestVI captures the expected pattern in atria, but it mistakenly treats cells related to cardiac neural crest as atrial cardiomyocytes. STdeconvolve successfully captures the spatial pattern, discerning even the difference between the ventricle and ventricular interval, but cell type mapping remains a problem for this reference-free method. Other methods tend to generate noisy cell type proportion estimates under this limited condition (**Supplementary Figure 4**). We further evaluate the performance when we let STdeconvolve choose its optimal cell type number in the same developing human heart slide (**Figure 2E-F**). In this evaluation, we only employed the ground truth cell types that can be successfully mapped to one STdeconvolve inferred cell type with a correlation greater than 0.5 (**Methods**). Comparing to mapping all the ground truth cell types, STdeconvolve shows improved performance in both RMSE and Pearson correlation evaluation (**Supplementary figure 5A-C**). STdeconvolve infers the number of cell types by minimizing the perplexity. Applying on the developing heart slide, STdeconvolve reaches the minimal perplexity at 13 cell types (**Supplementary figure 5C-D**). It is essential to have a prior knowledge about the number of cell types since STdeconvolve may fail to find the optimal cell type numbers within a given range. For example, applying to olfactory bulb slides using the default gene set, the perplexity keeps decreasing as the deconvolved cell types increase from 5 to 20, with 12 out of 20 cell types that have mean proportion smaller than 5% which we consider as an over-fitting result (**Supplementary figure 5E**).

### Evaluation with spot level resolution ST data

As a touchstone of the methods in realistic spot level ST data, we used a mouse brain dataset [18], an extensively studied and well-structured tissue. The mouse brain dataset was created through the commercially available 10X Visium Spatial platform [4, 18]. Spatial capture technology empowers the Visium Spatial platform through the use of spatially barcoded mRNA-binding oligonucleotides, demonstrating high-quality RNA-sequencing data with maintained two-dimensional positional information [4]. A major barrier in downstream analysis of the 10X Visium data is the resolution of each spot. Compared to the pooled ST spots discussed above, it is not feasible to use RMSE as the assessment metric in spot-level ST datasets as the gold standard, the true cell type proportions, is unknown. An alternative way to evaluate the performance is to assess performance of clustering results derived from inferred cell type proportions. Despite the extensive intermixing among different cell types within each layer, the layered and segmented structure of the mouse cortex can be observed at several levels: anatomic, major cell type, and layer-specific gene markers. In this regard, comparing the clustering results (using the inferred cell proportions) with pathologist annotated cortex layer structure (treated as working truth) (**Figure 3A, Supplementary Figure 6-7**) along with checking the major cell types could be considered as a reflection of the deconvolution performance. We used K-means to cluster the spots and then quantified the similarity between inferred clusters and annotated clusters using the AMI metric (**Figure 3B**). A higher AMI indicates higher similarity and better performance. Only the primary somatosensory (SSP) cortex area was used in our assessment.

**Figure 3.**
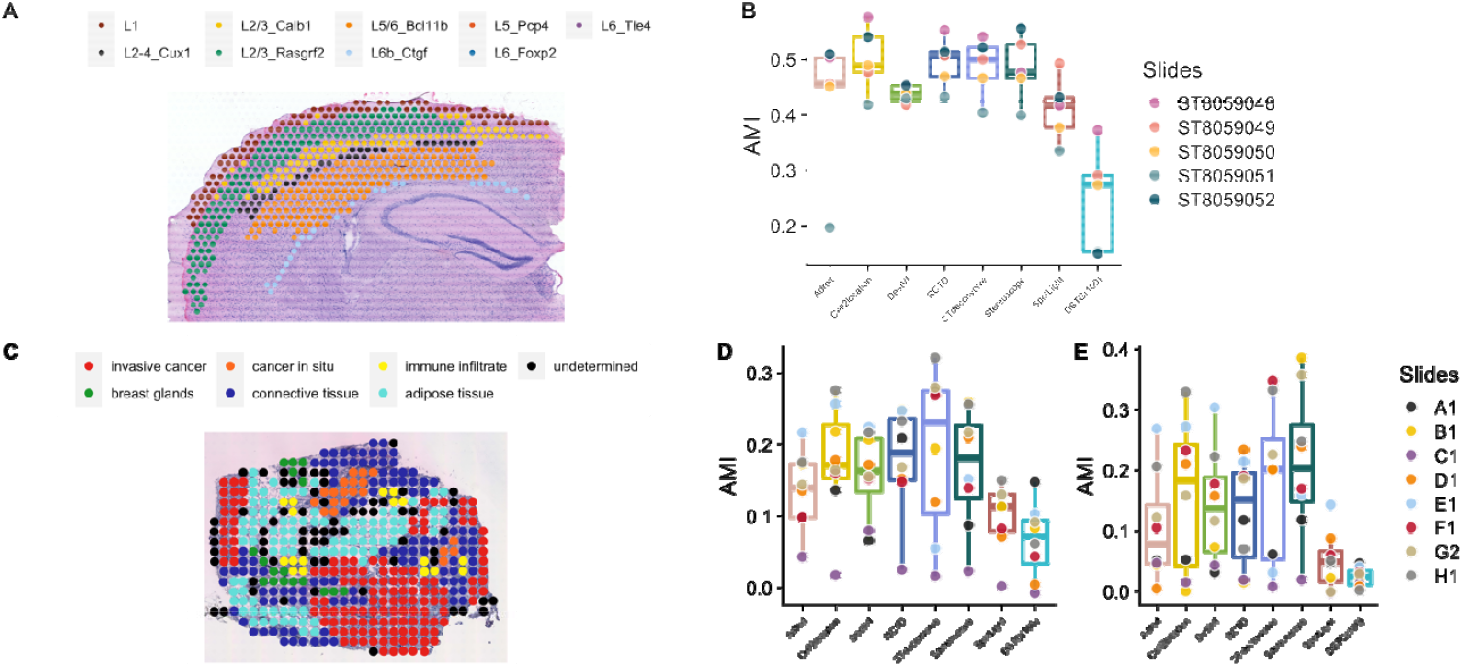
Evaluation on non-single cell resolution spot level ST data. (**A**) Histology image and pathologist annotation in the mouse brain data set (slide ST8059048). (**B**) AMI of K-means clustering results from 8 methods using the default gene subsets in the mouse brain tissue. (**C**) Histology image and pathologist annotation in breast cancer data set (slide G2). (**D**) AMI of K-means clustering results from 8 methods using the default gene subsets in the breast cancer data set. All clusters except for undetermined spots are used for evaluation. (**E**) AMI of K-means clustering results from 8 methods using the default gene subsets in the breast cancer data set. Clusters are combined to cancer and non-cancer areas. The cancer area contains both invasive cancer and cancer in situ areas. Other areas except for undetermined spots are combined as non-cancer areas. Then cancer and non-cancer binary classification is used in the evaluation. Undetermined spots are excluded in the evaluation.

Adroit, Cell2Location, RCTD, STdeconvolve, and Stereoscope all attain high AMI (**Figure 3B**). Clusters corresponding to L1 and L6b are relatively harder to identify due to the limited number of spots in these two layers (averaged 8.76% and 5.47% of the spots respectively over five slides). Cell2Location and STdeconvolve can effectively separate L1 from other layers, while DestVI, RCTD, SPOTlight, and Stereoscope successfully separate L6b from other layers. All methods except DSTG can recover the layered structure (**Supplementary Figure 7-12**). The expected major cell type distribution is observed in L2/3 IT CTX in the L2-3 layer, L4 IT CTX in the L4 layer, L4/5 IT in the L5 layer, and L6 CT CTX in the L6 and L6b layer (**Supplementary Figure 8-12**). The proportional estimates of cell types compared to the proportional estimates of cell types in each cortical layer of the Allen reference are remarkably consistent [18]. Adroit and Stereoscope suggest a smoothed-out dominating pattern (for example, L2/3 IT CTX-1 is the major cell type in L2/3), whereas Cell2Location and DestVI demonstrate a sharper dominating pattern. It appears that STdeconvolve obtains the top three AMI on four of five slides, in contrast to the previous results when applied to single-cell resolution ST data. One reason is that K-means does not require cell mapping. Note that STdeconvolve cannot be used to infer cell type mixtures for certain cell types. We cannot guarantee that the inferred cell types cover all the cell types we need, even when we set the number of cell types in STdeconvolve to our desired number, which can be considered as a drawback of the method. Despite these limitations, STdeconvolve is still able to correctly determine the layer structure. For example, in the slide ST8059049, inferred cell type 1 corresponds to L1 and cell type 11 corresponds to L6 (**Supplementary Figure 9**), thereby providing a much clearer pattern than most other methods.

Lastly, we demonstrate the performance on breast cancer [20], which is a less structured tissue (**Figure 3C**). There are several subtypes of breast cancer, among which the HER2-positive subtype is characterized by the increased expression of *HER2* (human epidermal growth factor receptor 2) in tumor cells [20]. We obtained ST data of HER2-positive tumors from eight individuals (patient A-H) generated through the 10X Visium Spatial platform [4, 20]. In the analysis, we had eight slides, one from each patient, all of which are pathologist-annotated, containing areas with one of the following five labels: in situ cancer, invasive cancer, adipose tissue, immune infiltrate, or connective tissue (**Supplementary Figure 13**) [20]. Further, we created another set of labels where invasive cancer and cancer in situ are combined as cancer area, while all other areas except for undetermined spots are grouped as the non-cancer area (**Supplementary Figure 14**).

AMI in the breast cancer tissue is substantially lower/worse for all the methods compared to the result in the mouse brain analysis presented above. As mentioned earlier, this is expected since these tumor samples tend to be more complex and much less structured than the mouse cortex. The B cells, T cells, and Plasmablasts cells are tightly embedded in the tSNE embedding space. Normal epithelial cells and cancerous epithelial cells appear to differ by some distance. This will be the main focus during the assessment, along with the annotated cancer regions created by the pathologist (**Supplementary Figure 2**).

DSTG remains as the worst performer with the lowest AMI when assessed with either set of labels, namely the five-category set or the binary set separating only cancer vs non-cancer areas. The best performers with highest AMIs are Cell2Location, RCTD, Stereoscope, DestVI, and STdeconvolve. When evaluating using the five-category labeling set, the AMI of STdeconvolve varies widely across 8 slides, while other methods exhibit a more consistent pattern (**Figure 3D**). When using the binary (cancer vs non-cancer) labeling in evaluation, AMI varies widely in most of the methods (**Figure 3E**). In terms of separating cancer from non-cancer areas, Stereoscope appears to be the most effective with the highest average AMI across all methods evaluated (**Figure 3E**). Although all the methods have a relatively low AMI, a few of them manage to reveal expected cell type enrichment in their corresponding areas (**Supplementary Figure 15-22**). For example, Adroit, Cell2Location, RCTD and Stereoscope consistently show enrichment of cancer epithelial cells and depletion of normal epithelial cells in cancer areas. Although DestVI fails to capture these expected patterns in a few patients (particularly patient A1, F1, G2 in **Supplementary Figure 15,20-21**), the cancer epithelial mapping shows the clearest boundary between cancer and non-cancer areas in other patients (patient B1, D1, **Supplementary Figure 16,18**).

As a conclusion, Adroit, Cell2Location, RCTD, DSTG, and Stereoscope all achieve excellent performance when the single cell reference and the ST dataset are matched. When only a small number of genes are available, Adroit, Cell2Location, RCTD, and Stereoscope are still effective in inferring the cell type proportion. Clustering based on inferred cell proportions demonstrates strong agreement with pathologist-assigned layers for all methods except for DSTG in well-structured tissues. In the case of less-structured tissues, all methods struggle to infer a spatial pattern in agreement with gold standard annotation. Overall, Adroit, Cell2Location, RCTD, and Stereoscope outperform others. While DestVI performs well on some of the samples, it is less robust compared to the four consistent outperformers.

## Discussion

As ST technologies continue to evolve rapidly, we anticipate having more advanced technology that captures single-cell resolution data as well as measures expression levels of as many genes as possible. Currently, spot-level resolution ST data still has its advantages in the whole transcriptomics-wide gene measuring capability. On the data analysis side, inferring composition of cell types has already been demonstrated to be helpful in downstream analysis such as detecting spatial expression patterns in spot-level data [25]. Such downstream analysis is essential both for technical reasons (e.g., identifying spatial domains and spatially variable genes [26] in the gene selection process when designing experiments generating single-cell resolution ST data) and for revealing novel biological insights without suffering from impaired power or inflated false positives. To perform an appropriate downstream analysis, it is critical to perform cell type mixture decomposition in ST spots. An incorrect inference of cell mapping could lead to misunderstandings of tissue structure and function. In this review, we have benchmarked the performance of eight ST deconvolution methods using four real datasets. This work can be decomposed into two main parts.

First, our study evaluates the performance in two single-cell resolution datasets from mouse olfactory bulb [3] and developing human heart [19], respectively. We pool the cells according to their spatial coordinates so that a gold standard is established using the true cell type proportions based on the contributing cells at each constructed spot. We quantify the deconvolution performance using RMSE and Pearson correlation. We further examine the proportion of each cell type. Most ST deconvolution methods claim to maintain their performance even when there are systematic differences between the single cell reference and the ST data under inference [9, 11, 18]. For example, Stereoscope [11] claims to be robust to missing cells. Both Stereoscope and cell2location are reported to well accommodate technology differences in measuring gene expression [11, 18]. When we evaluate the methods in those single-cell resolution datasets, the single cell reference and ST data are perfectly matched as the same single cell resolution data serve as the reference and are used to construct ST spot. Thus, the results do not reflect the ability of these methods to deal with inconsistency between reference and ST data, but rather indicate whether their assumptions ruin their performance when there is a perfectly matching reference. Moreover, with several different gene sets, we have benchmarked the robustness and time complexity, providing insights regarding the best choice of gene subset for each method. Furthermore, as the number of genes available in the heart data is limited, these methods are further examined regarding their ability to deconvolve when the number of available genes is substantially reduced.

In the second part of the analysis, we examine the methods on two spot-level resolution ST datasets, including mouse brain [18] and HER2+ breast cancer [20]. The true cell type proportion, also known as the gold standard, is unknown for spot level data. The mouse brain, as a tissue that has been extensively studied, is a valuable example to start with since it retains a clear structure for pathologists to annotate. We can assess the deconvolution results based on known molecular, anatomical, and functional information that are well established with previous studies. Specifically, using the cell type proportion estimates, we cluster the spots. The K-means-inferred clusters as well as the major cell type in each spot are further compared with their pathologist-annotated layers. Following this, the performance is assessed on a much less structured tissue from breast cancer samples, in which we test the ability to distinguish cancer regions using clustering results (again via K-means clustering from inferred cell type proportions) and examine whether the proportion of cancerous cells mirrors the cancerous areas annotated by pathologists.

In summary, there is no method that consistently outperforms others across tissue types. Most of the probabilistic methods tested in our study, especially Adroit, RCTD, Cell2Location, and Stereoscope exhibit consistently high performance across tissues. STdeconvolve, as the only reference-free method, has the capability for identifying tissue structure and cell mixture, but the cell type mapping must be addressed carefully. We have thoroughly evaluated various scenarios, encompassing different tissues, varying technologies and data resolution, different numbers of single cells and spots, as well as number and type of genes employed for analysis. We therefore recommend that investigators first identify some of our evaluated scenario(s) that best match their own data and select best performing methods under these scenario(s). With the use of the appropriately chosen methods and gene sets, we hope the increased accuracy of cell mapping inference will assist in the future downstream analyses. In addition, while out of the scope of this work, denoising and dimension reduction of noisy and high dimensional ST data can allow more effective information extraction [27]. We anticipate that cell type deconvolution further benefits from development and advancement of methods that effectively denoise and reduce the dimension of ST data.

## Data availability

All datasets used in this study are publicly available. The detailed reference and the websites to download them are in Table 2. The pathologist’s annotation of five mouse brain slides is available upon request.

## Acknowledgements

We thank Li lab members for providing advice on data preprocessing, method selection and feedback on the manuscript. Figure1 is created via bioRender.

## Financial support

The research is partly supported by NIH grants U01HG011720, U01DA052713, and P50HD103573.

## Conflicts of interest

The authors declare that they have no conflict of interest.

## Methods

### Data preprocessing

We utilized four data sets to test the ST deconvolution methods in this study. The data preprocessing steps are summarized as follows.

SeqFISH+ is a single-cell level ST data. For each field of view (FOV), cells that locate in the same 400×400 square pixel area are pooled as a spot. The single-cell level data is also used as the single-cell reference. Genes expressed in at least 3 cells and cells that have at least 200 features and at most 2500 features are kept in the single-cell reference.

The human heart ST data is also a single-cell resolution ST data. We analyzed the PCW6.5_1 slide. Genes expressed in at least 1 cell and cells that have gene expression greater than 0 are kept in the single-cell reference. Cells that are co-located in the same 454×424 square pixel area are pooled as a ST spot.

ST data from mouse brain primary somatosensory region (SSp) are obtained from (the cell2location paper). Mouse brain single-cell data was obtained from Yao et al (2021) and only cells from the SSp region were kept in the analysis as reference. For ST data, we only kept genes that are present in at least 2% of spots. For scRNA-seq data, we removed cells that were not confidently assigned a class label by the original paper. Only genes that exist in both scRNA-seq and ST data are used in the following analysis.

In the breast cancer analysis, we only keep HER2+ subtype ST data for the deconvolution evaluation as well as keeping only the HER2+ patients’ scRNA-seq data as reference. For scRNA-seq reference, genes expressed in at least 3 cells and cells expressing at least 200 genes are kept. Similarly for ST data, genes expressed in at least 3 spots and spots expressing at least 100 features are kept. Only genes that exist in both scRNA-seq and ST data are employed in the further analysis.

### Gene subsetting and gene set employed in the ST deconvolution methods

All gene subsettings are accomplished using the R package Seurat [28]. Highly variable genes are selected using feature variance calculated by the FindVariableFeatures function with default settings. Marker genes are selected using the FindAllMarkers function with a log-fold-change threshold of 0.75. Both positive and negative markers are included in the marker gene set. Top marker genes are selected using p values.

### Default gene set used in each deconvolution method

Adroit, Cell2Location, DestVI, Stereoscope [10, 11, 16, 18] do not have a built-in gene filtering strategy. Top 2000 highly variable genes are used in the analysis. SPOTlight [15] used a mixture of top 500 marker genes for each cell type and the top 500 highly variable genes. Note here we follow the pipeline and keep only positive markers when selecting marker genes. We used the default parameters of RCTD and ran RCTD in full mode. By default, RCTD [9] has a built-in marker gene selection step where only genes with normalized gene expression greater than or equal to 0.0002 are included, and it selects cell type marker genes based on a log-fold-change threshold of 0.75. Only selected cell type marker genes are input to the algorithm. During STdeconvolve inference, following the software pipeline, we removed genes detected in less than 2% of spots or genes that are expressed in all spots. STdeconvolve then selects genes by choosing significantly over-dispersed genes across spots to detect transcriptionally distinct cell types. By default, only the top 1,000 or fewer most over-dispersed genes are retained in the input ST data. DSTG selects the top 2,000 most variable genes by default across different cell types in the reference scRNA-seq data according to adjusted P-values with Bonferroni correction using the analysis of variance.

Note that after selecting the gene subset, there may appear a situation that some of the spots do not have the selected gene(s) expressed. Such spots are removed from further analysis.

### Other Technical details

In the olfactory bulb dataset, 7 fields of views are combined as one input file for all 8 methods. For the other 3 tissues, the inference and deconvolution were performed separately on each single slide.

STdeconvolve additionally removes spots with library size smaller than 100. We followed this guidance for development human heart, mouse brain and breast cancer slides. But for the SeqFISH+ data, we chose to be more lenient and kept all spots with library size > 0 since the spot number is limited (only 164 spots total). Moreover, STdeconvolve filters out genes expressed in fewer than 2% of the spots in all datasets. Finally, we set the number of clusters the same as the number of cell types in the corresponding reference scRNA-seq dataset. All other parameters were set as default.

DSTG has a default setting of 2,000 HVGs and does not run when users input data with fewer than 2,000 genes. We set k=100 as the number of neighbors in our analysis in the mouse brain and breast cancer analysis. In the olfactory bulb data analysis, we used k=20, 50, 100. All other parameters were set as default.

We used 6,000 epochs in the single-cell inference and 30,000 epochs in the ST deconvolution for Cell2Location. We used 500 epochs in the single-cell inference and 4,000 epochs in the ST deconvolution for DestVI. For Stereoscope, 30,000 epochs were used in both single-cell inference and ST deconvolution. For SPOTlight, 300 cells per cell type were employed in the analysis.

To annotate cell types identified by STdeconvolve, we used the transcriptional correlations method described in the software, where we computed the Pearson’s correlation between every combination of deconvolved and ground truth cell-type transcriptional profiles, and matched each deconvolved cell type with the ground truth cell-type with the highest Pearson’s correlation. We only mapped one ground truth celltype to each deconvolved cell type. We used the same scRNA-seq dataset as in other methods for the ground truth reference. For the evaluation when we let STdeconvolve choose the optimal number of cell types, we only employed the ground truth cell types that can be successfully mapped to one STdeconvolve inferred cell type with a correlation greater than 0.5. The “successfully” inferred cell types are defined as ground truth cell types that have a large than 0.5 correlation with deconvolved cell types. Note here, there may be some ground truth cell types that have larger than 0.5 correlation with deconvolved cell types but are still excluded from the evaluation. This is because that the corresponding cell type have larger correlation with other ground truth cell types. For example, for the developing human heart tissue dataset (**Supplementary Figure 5c**), ground truth cell type 4 has 0.88 correlation with deconvolved cluster 1 and 0.87 correlation with deconvolved cluster 4. However, deconvolved cluster 1 has 0.94 correlation with ground truth cell type 2 and deconvolved cluster 4 has 0.93 correlation with ground truth cell type 3. Besides deconvolved clusters 1 and 4, the highest correlation for ground truth cell type 4 is 0.44. With the above conditions, ground truth cell type 4 is excluded from the evaluation.

T-SNE coordinates were calculated using Seurat_3.2.3 in R v3.6.0. We analyzed each scRNA-seq dataset following the standard pipeline with default Seurat parameter setting. Principle component analysis (PCA) was performed on genes using RunPCA and the top 10 PCs were used as input to both Rtsne method using RunTSNE in Seurat.

## Notes

### Competing Interest Statement

The authors have declared no competing interest.

