## Supplementary information for "A comprehensive comparison on cell type composition inference for spatial transcriptomics data"

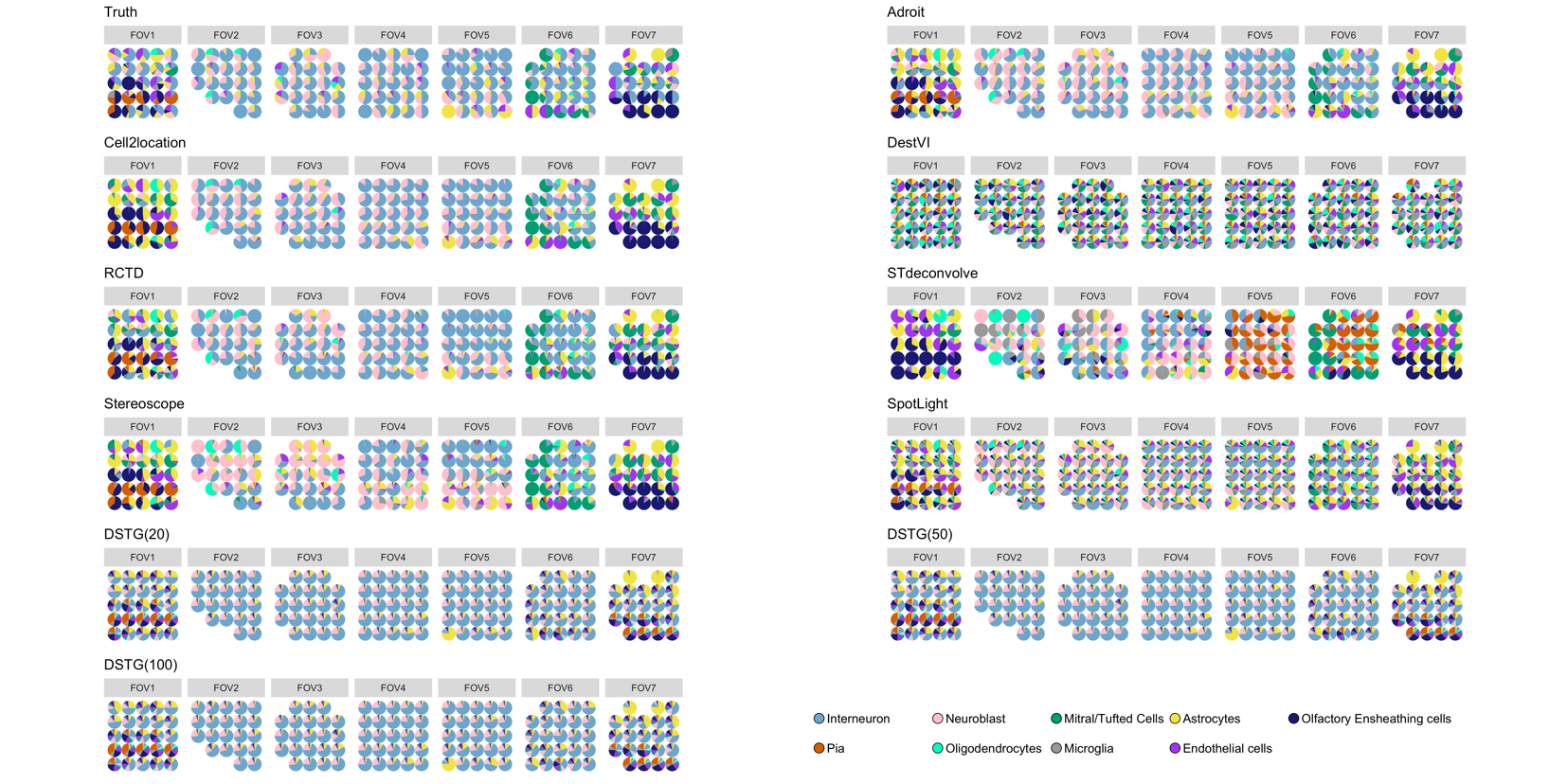

Supplementary Figure 1. Cell type proportion estimates in mouse olfactory bulb data. Default gene set is used here. Pie chart at each spot shows the proportions of estimated cell types for the corresponding spot.

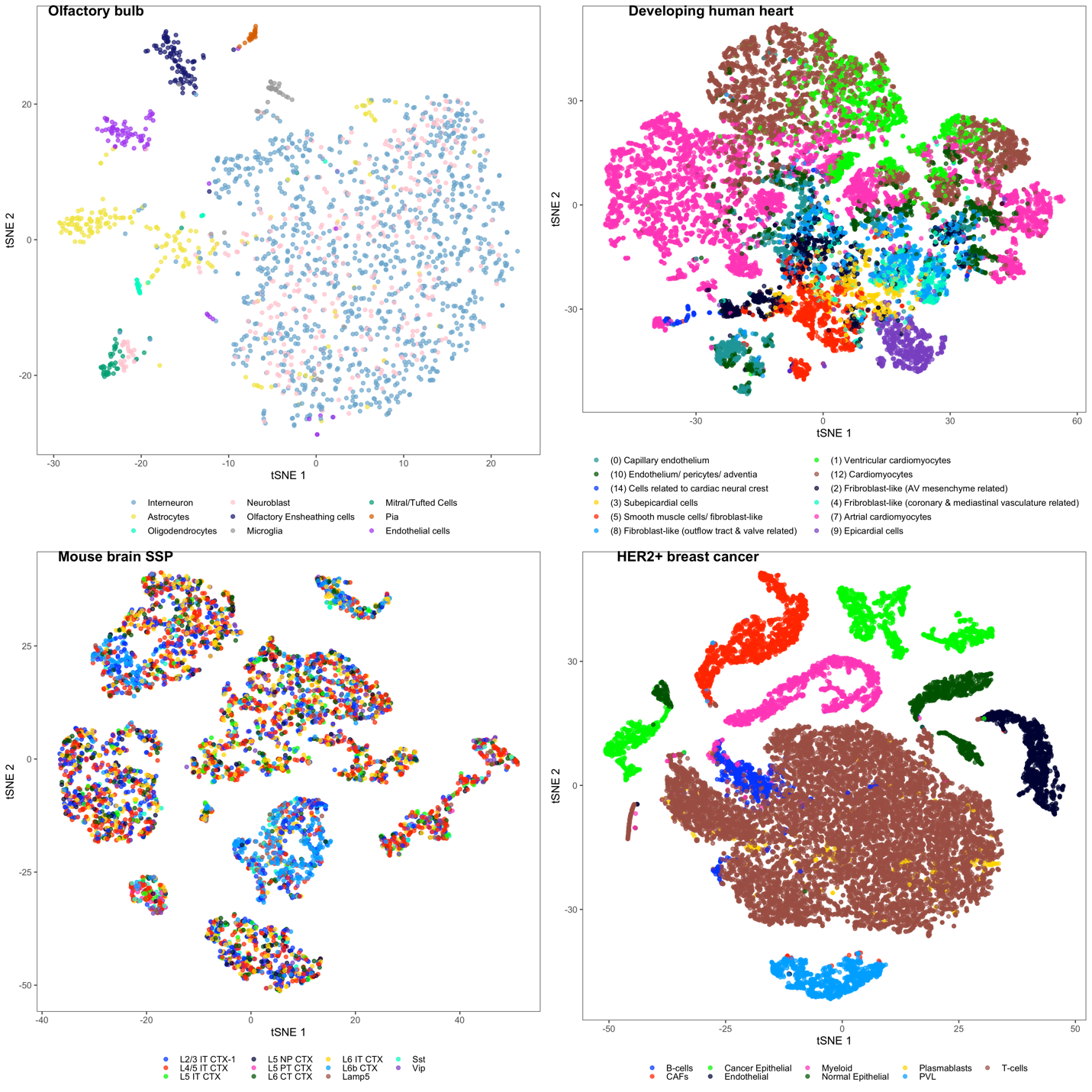

Supplementary Figure 2. tSNE embedding of cells from the four scRNA-seq reference data.

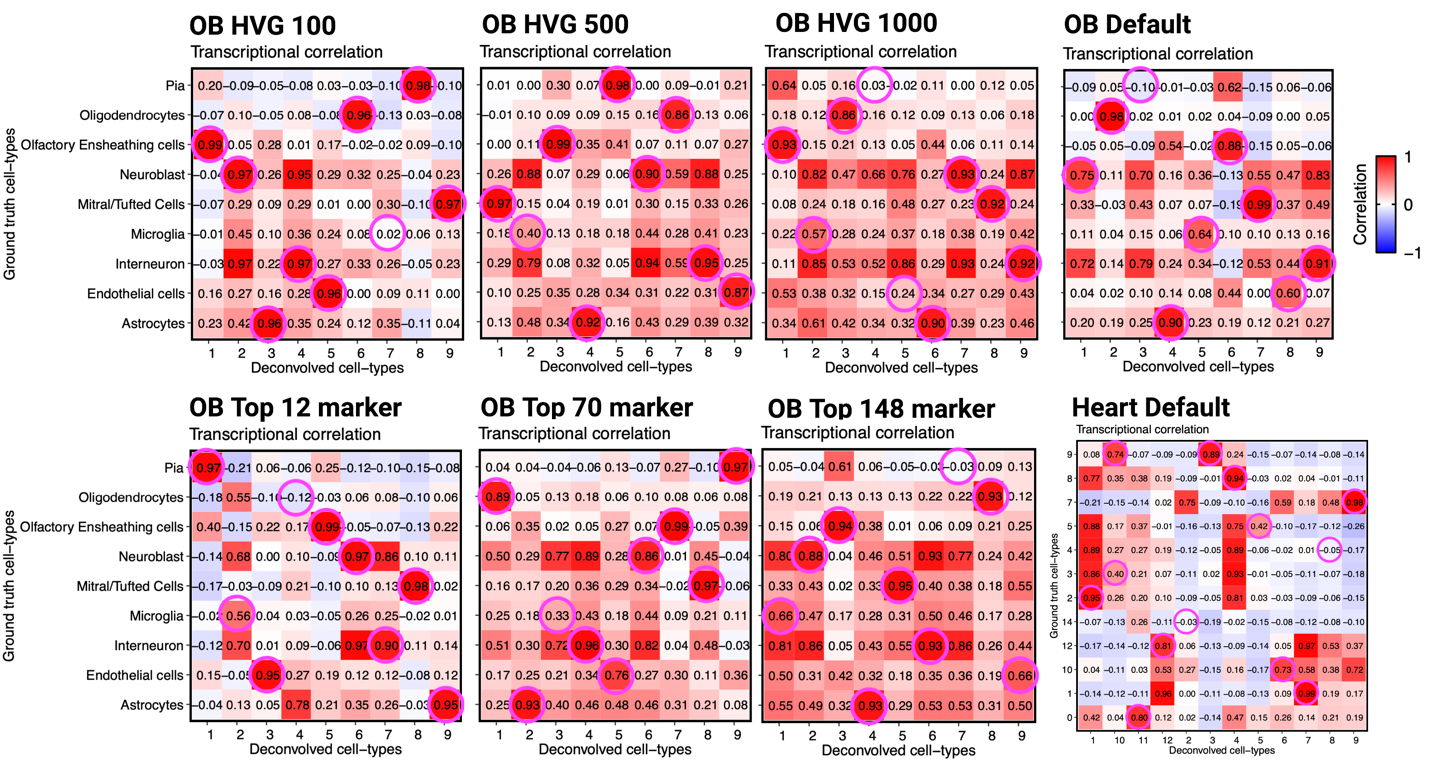

Supplementary Figure 3. Cell type mapping for STdeconvolve. OB: olfactory bulb data. Heart: developing human heart data. The pink circle indicates the cell type assignment used in the evaluation. The ground truth cell types for developing human heart tissue are labeled identically as in Figure 2.

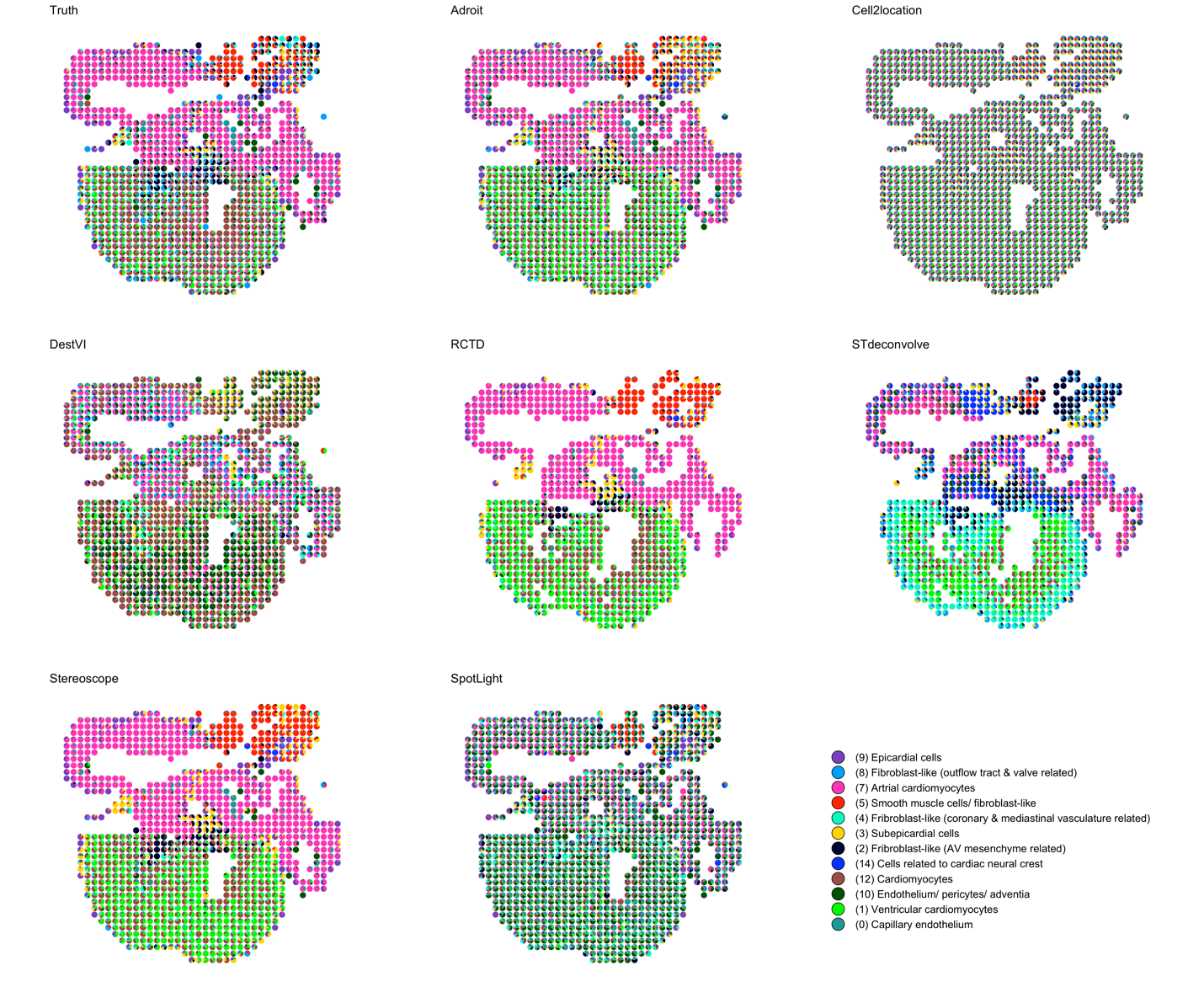

Supplementary Figure 4. Cell type proportion estimates in developing human heart tissue. If a method has no built-in gene selecting strategy, all genes are used for inference.

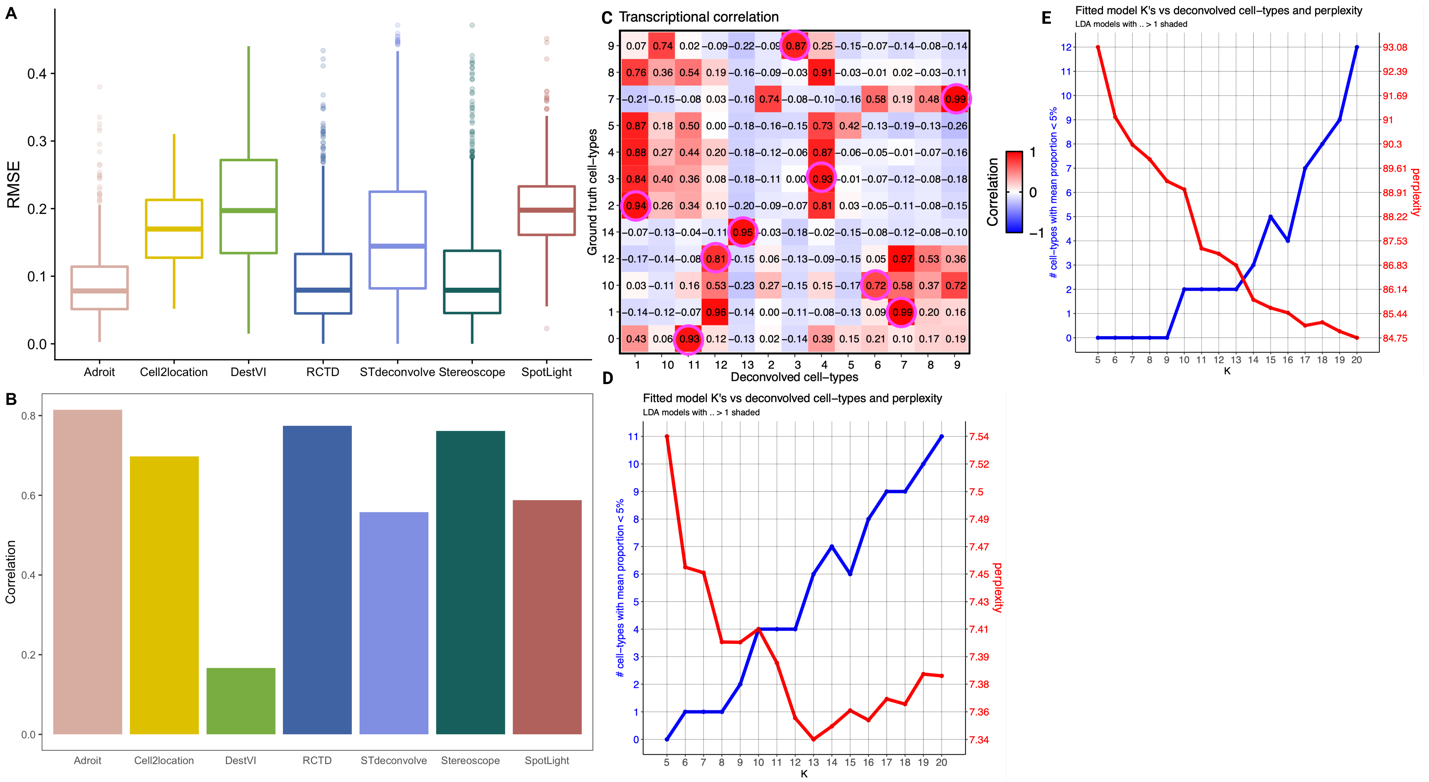

Supplementary Figure 5. Evaluation with only cell types that STdeconvolve “successfully” inferred. (**A**) RMSE of heart cell proportion estimates from 8 methods using the default gene set for the human heart dataset. (**B**) Pearson correlation of heart cell proportion estimates from 8 methods using the same sets of genes as in **A**. (**C**) Cell type mapping for STdeconvolve in the developing human heart tissue. The pink circle indicates the cell type assignment used in the evaluation. The ground truth cell types are labeled identically as in Figure 2. (**D**) Optimal cell type number selecting process employed by STdeconvolve in the development human heart dataset with default gene set (**E**) Optimal cell type number selecting process employed by STdeconvolve in the olfactory bulb data set with default gene set. K with the lowest perplexity is treated as the optimal cell type number.

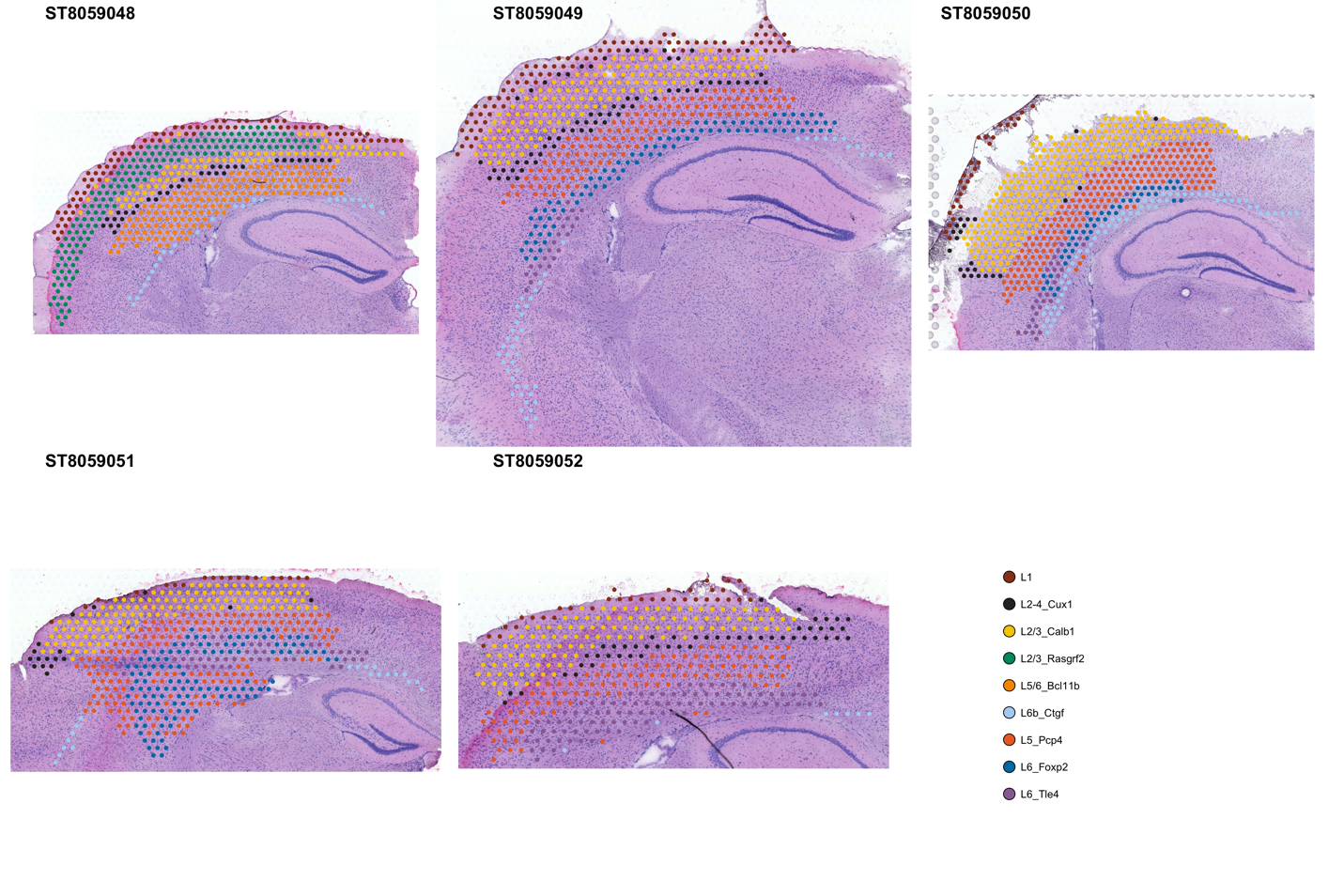

Supplementary Figure 6. Pathologist annotation for 5 mouse brain slides.

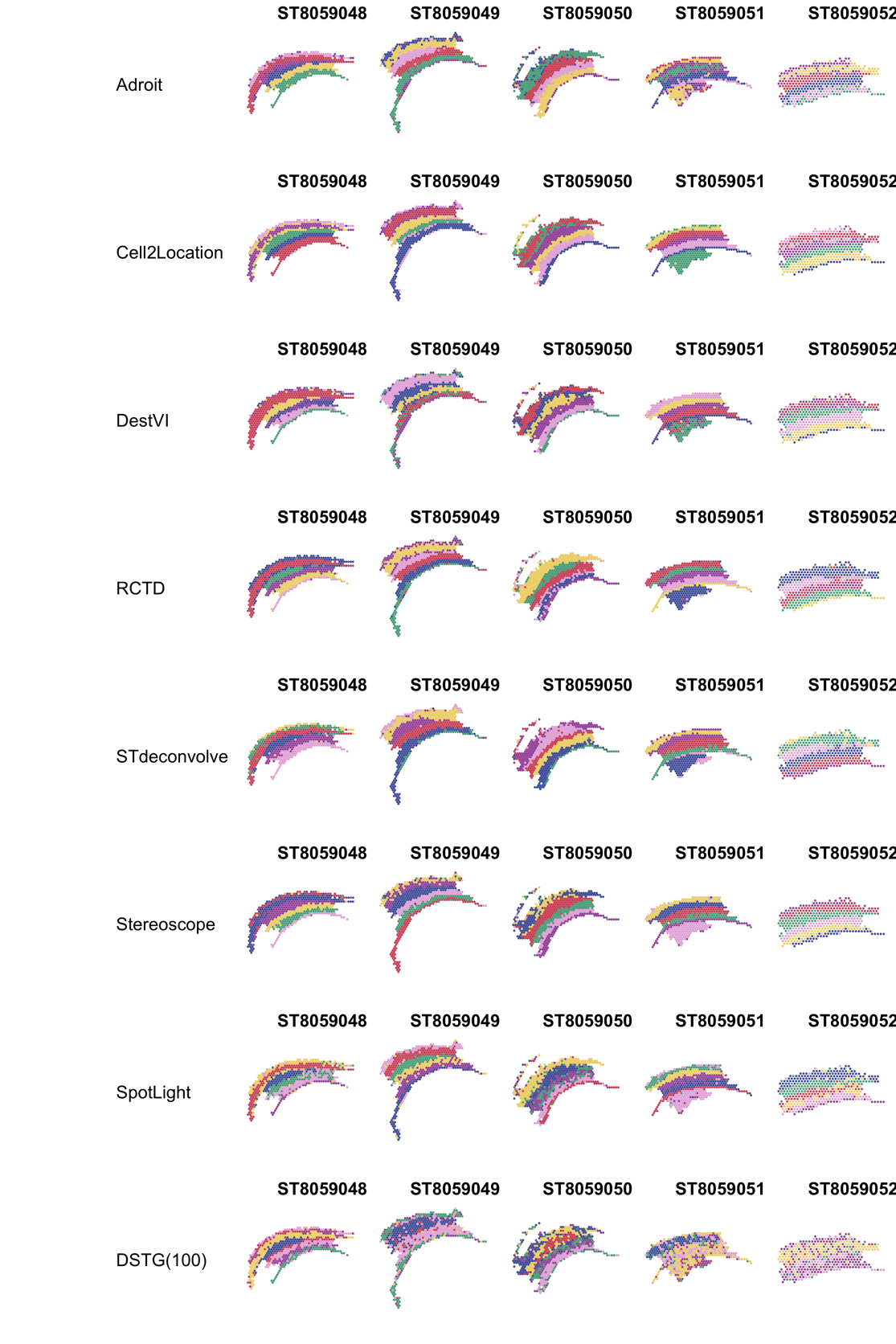

Supplementary Figure 7. K-means inferred clusters (n=6) in the mouse brain tissue. The colors of each spot indicate the inferred cluster.

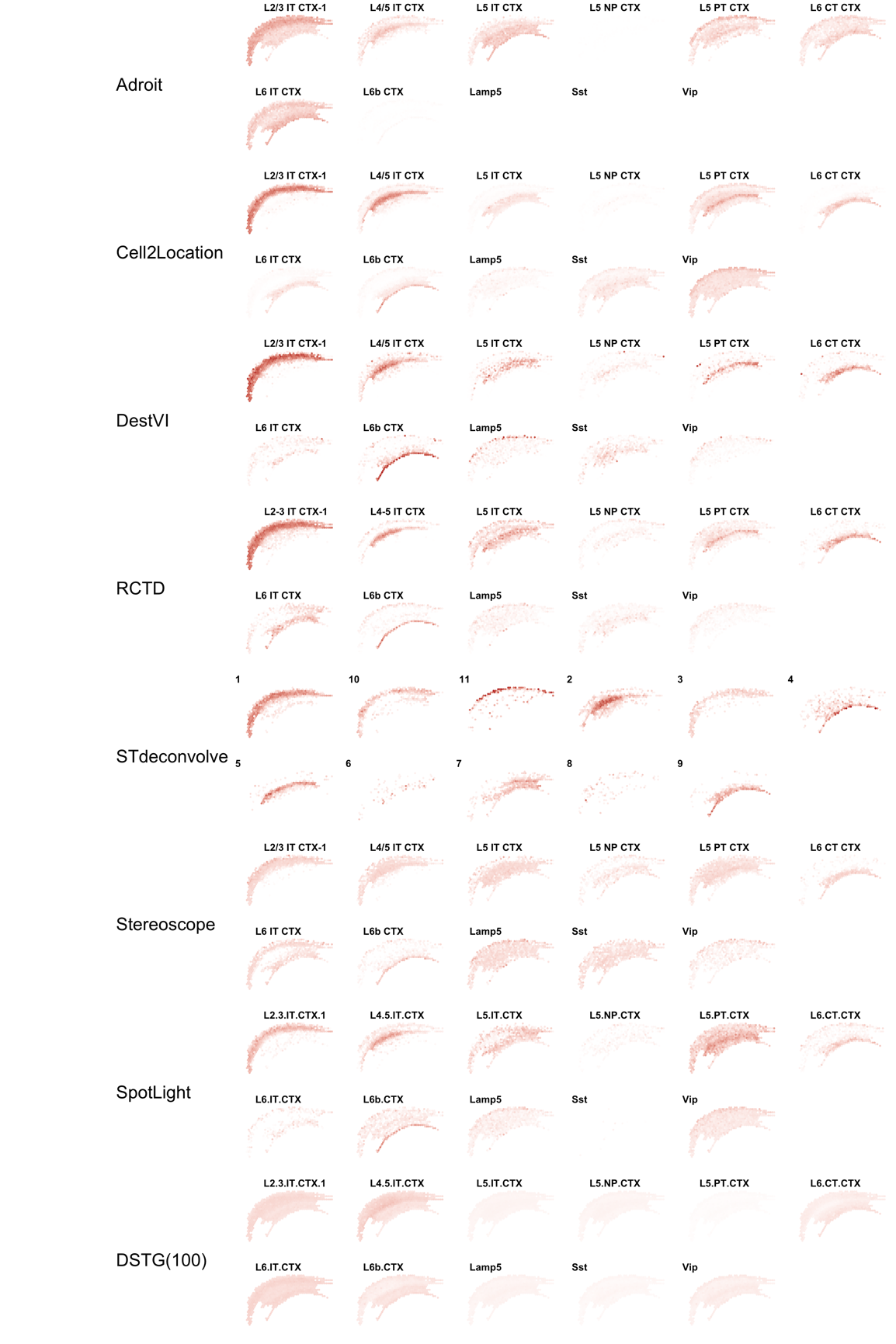

Supplementary Figure 8. Normalized cell type proportion estimates of the ST8059048 slide. Cell type proportion estimates are normalized to sum up to 1. White indicates 0 and red indicates 1 such that the redder the color, the closer to a proportion of 1.

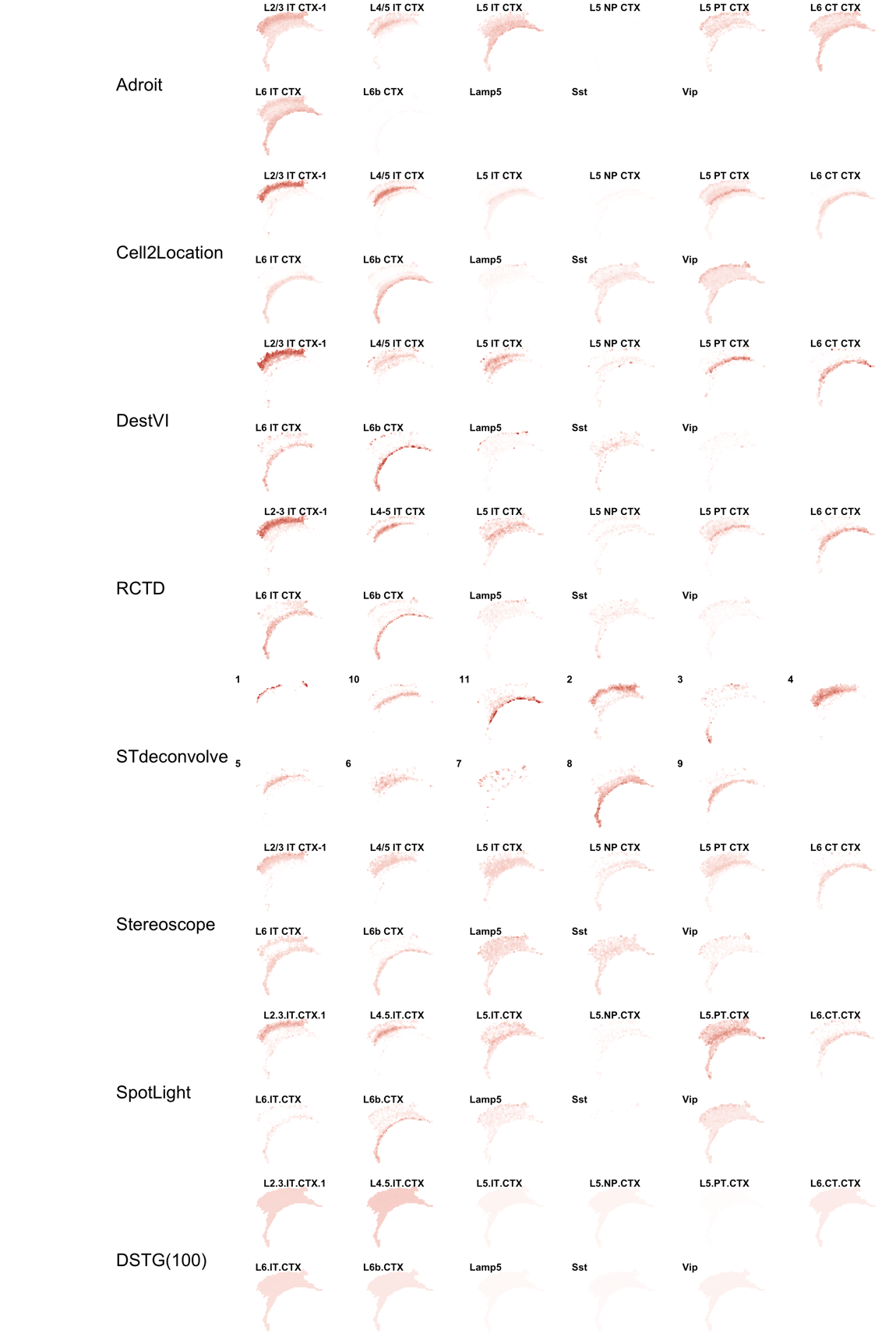

Supplementary Figure 9. Normalized cell type proportion estimates of the ST8059049 slide. Cell type proportion estimates are normalized, to sum up to 1. White indicates 0 and red indicates 1 such that the redder the color, the closer to a proportion of 1.

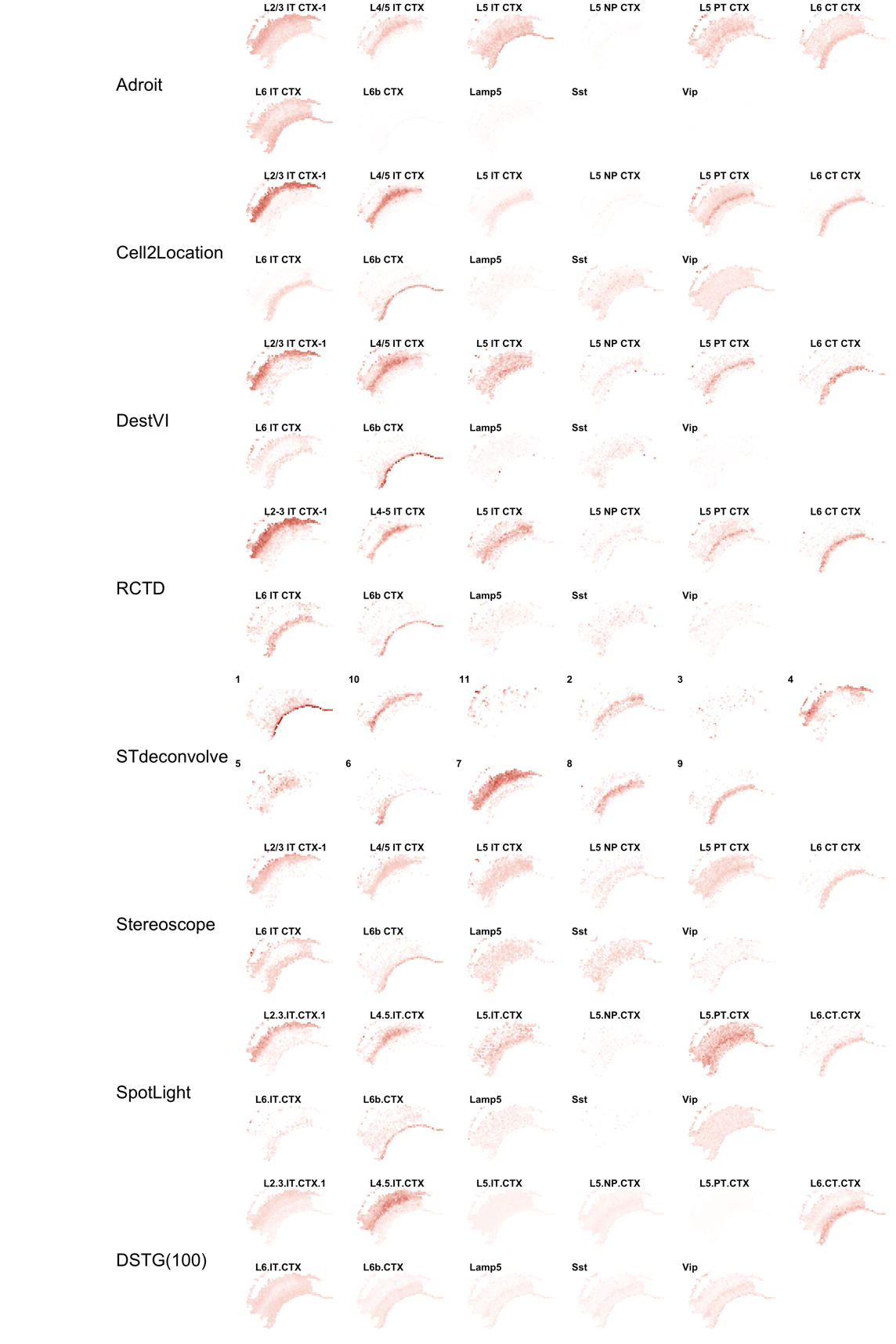

Supplementary Figure 10. Normalized cell type proportion estimates of the ST8059050 slides. Cell type proportion estimates are normalized, to sum up to 1. White indicates 0 and red indicates 1 such that the redder the color, the closer to a proportion of 1.

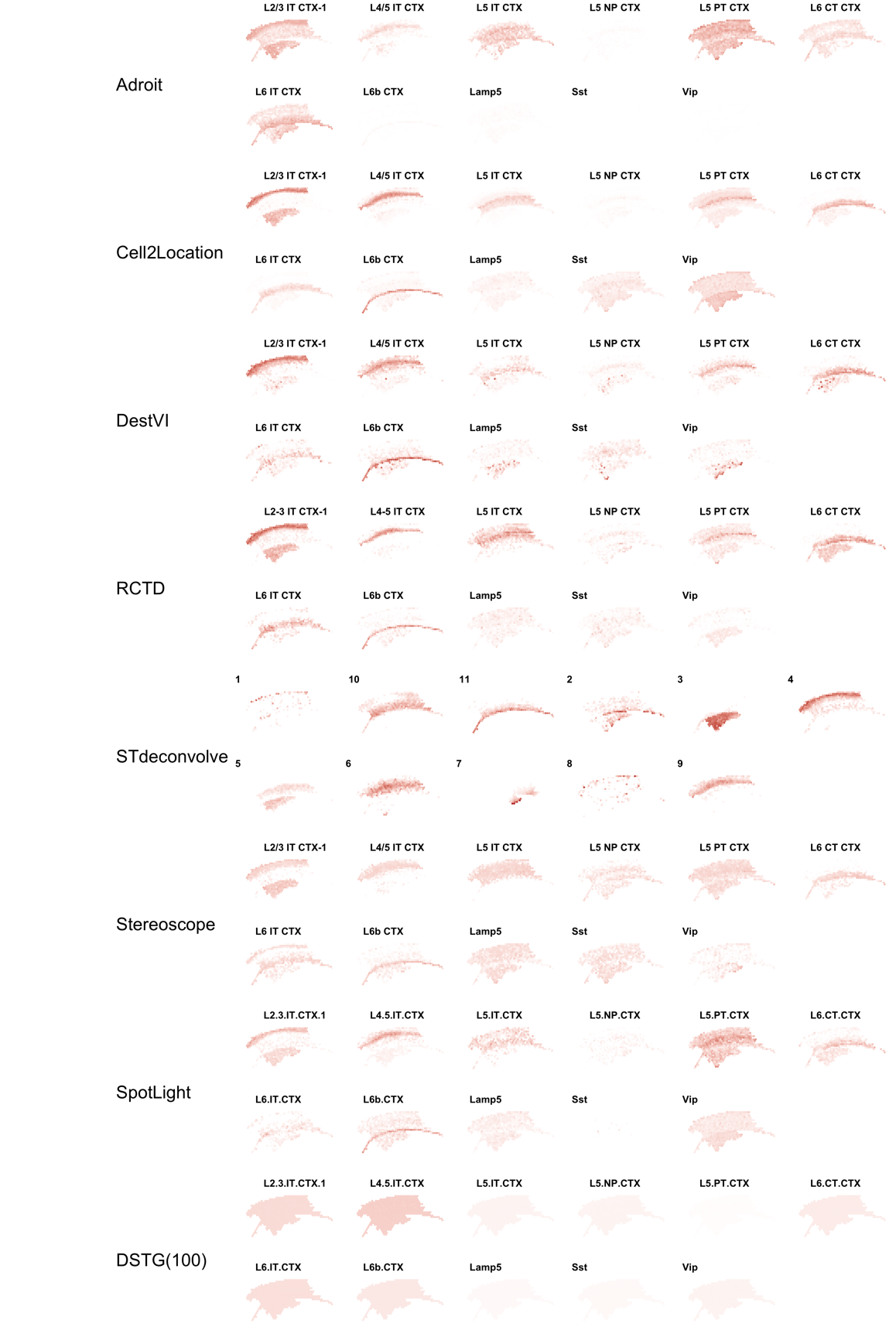

Supplementary Figure 11. Normalized cell type proportion estimates of the ST8059051 slide. Cell type proportion estimates are normalized, to sum up to 1. White indicates 0 and red indicates 1 such that the redder the color, the closer to a proportion of 1.

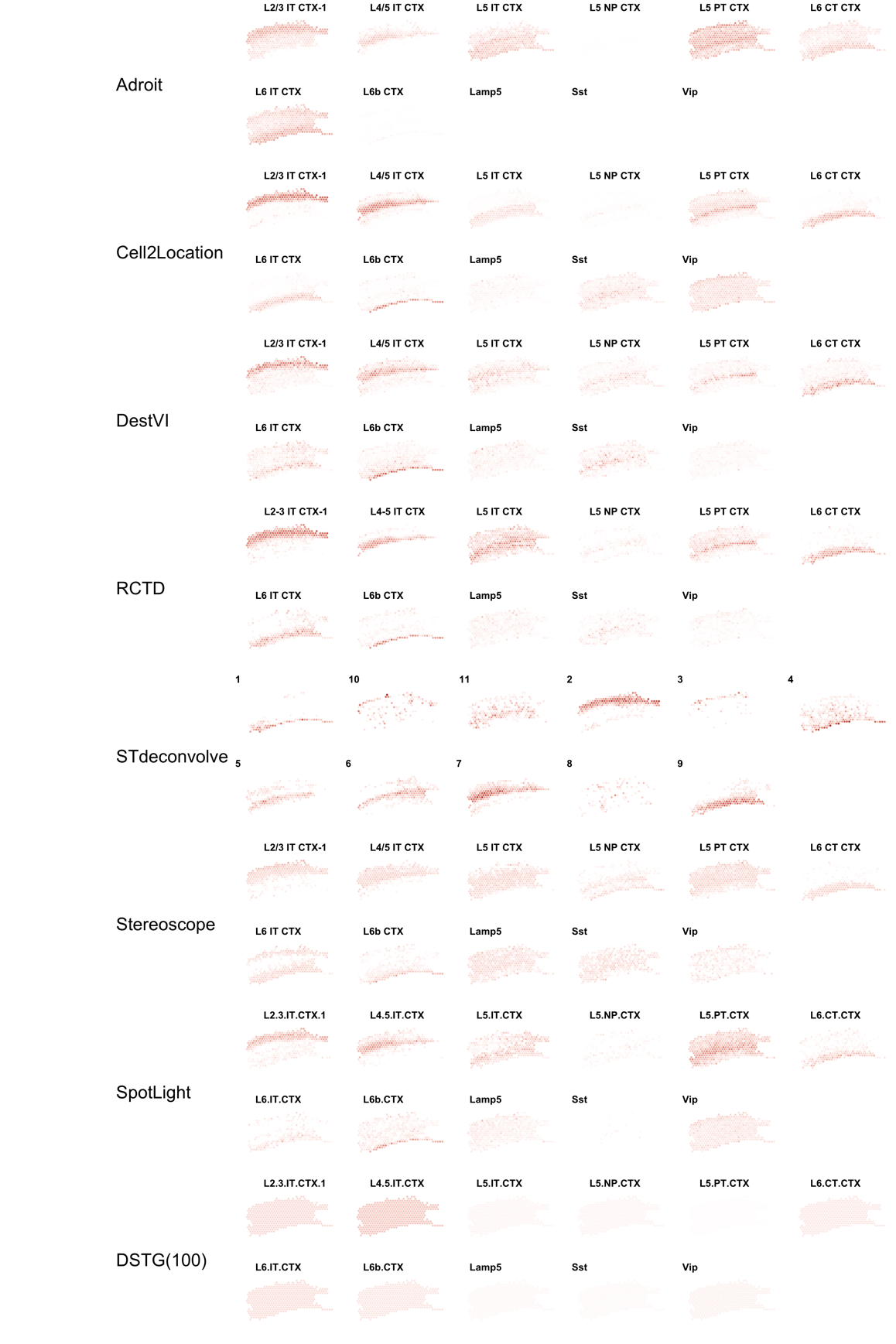

Supplementary Figure 12. Normalized cell type proportion estimates of the ST8059052 slide. Cell type proportion estimates are normalized, to sum up to 1. White indicates 0 and red indicates 1 such that the redder the color, the closer to a proportion of 1.

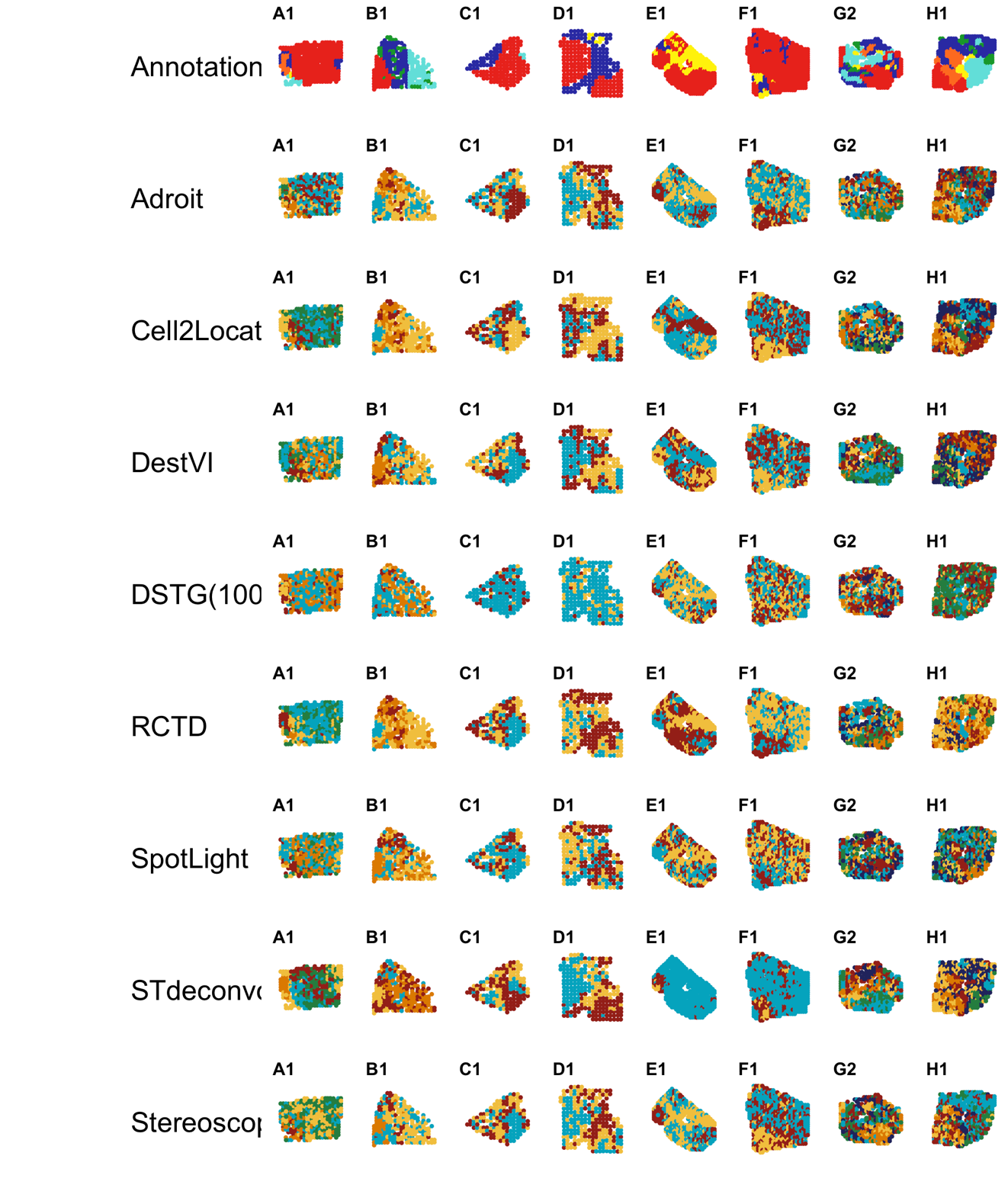
Supplementary Figure 13. K-means inferred clusters in the mouse brain tissue. The number of clusters for each slide used in the K-means inference is according to the number of annotated clusters in the annotation file for each layer. The first row is the pathologist's annotated clusters, in which the color coding follows that in Figure 3C. For rows 2-9, the colors of each spot indicate the inferred cluster.

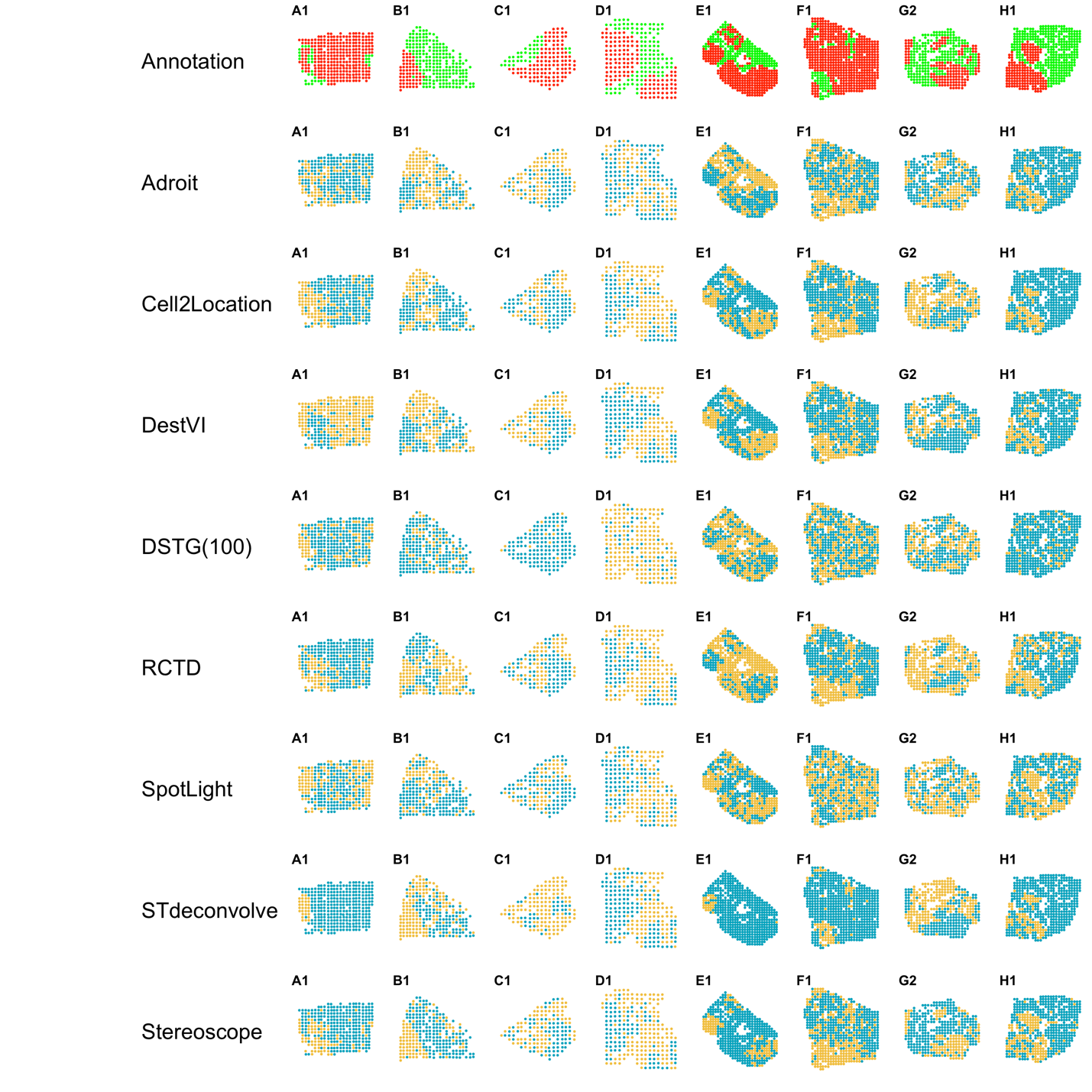

Supplementary Figure 14. K-means inferred clusters in the mouse brain tissue. Two clusters are used in the K-means inference. The first row is the pathologist's annotated clusters which are combined into cancer(red) and non-cancer areas (green). For rows 2-9, the colors of each spot indicate the inferred cluster..

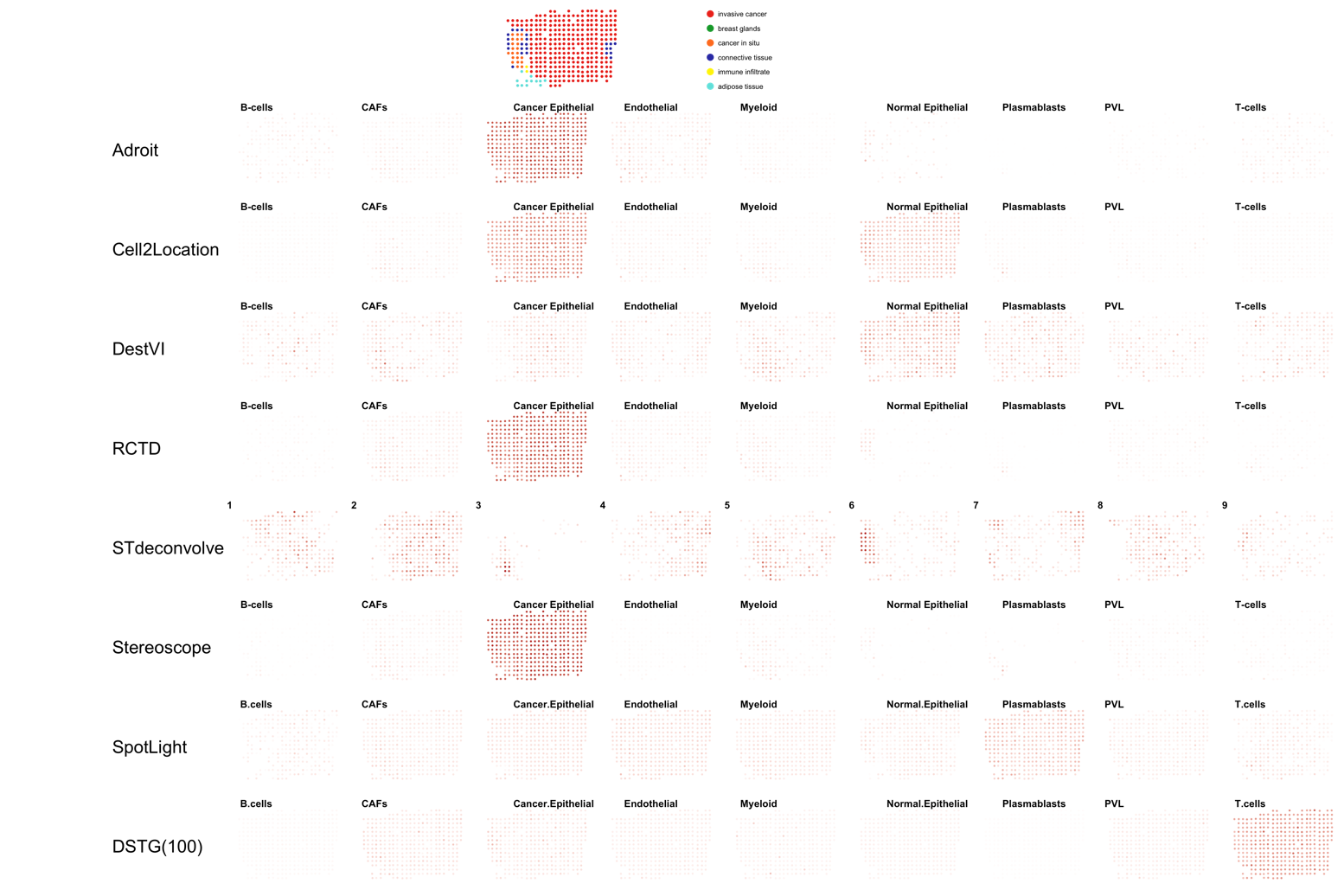

Supplementary Figure 15. Normalized cell type proportion estimates of the A1 slide. The first row is the pathologist annotated layer. Cell type proportion estimates are normalized, to sum up to 1. White indicates 0 and red indicates 1 such that the redder the color, the closer to a proportion of 1.

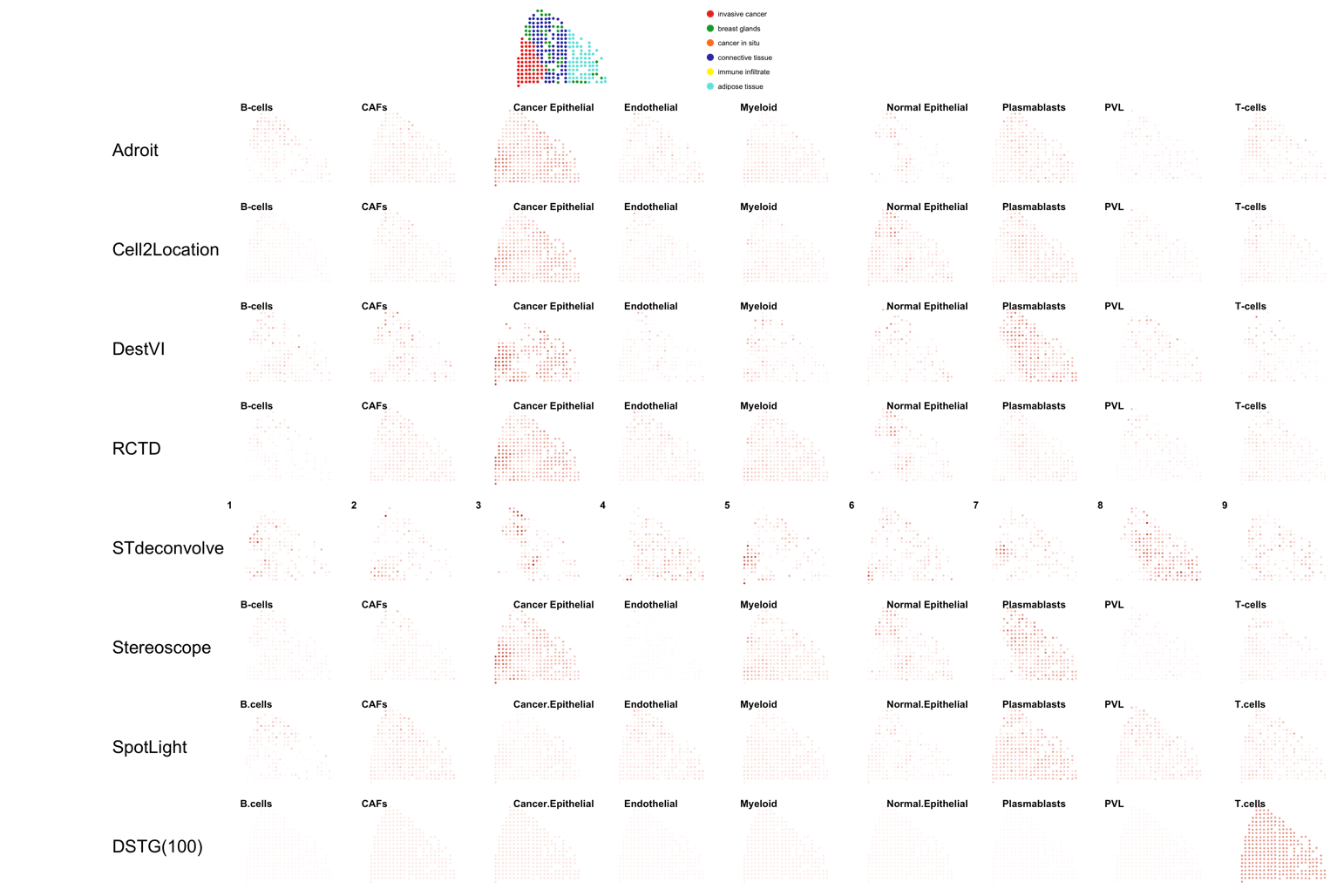

Supplementary Figure 16. Normalized cell type proportion estimates of the B1 slide. The first row is the pathologist annotated layer. Cell type proportion estimates are normalized, to sum up to 1. White indicates 0 and red indicates 1 such that the redder the color, the closer to a proportion of 1.

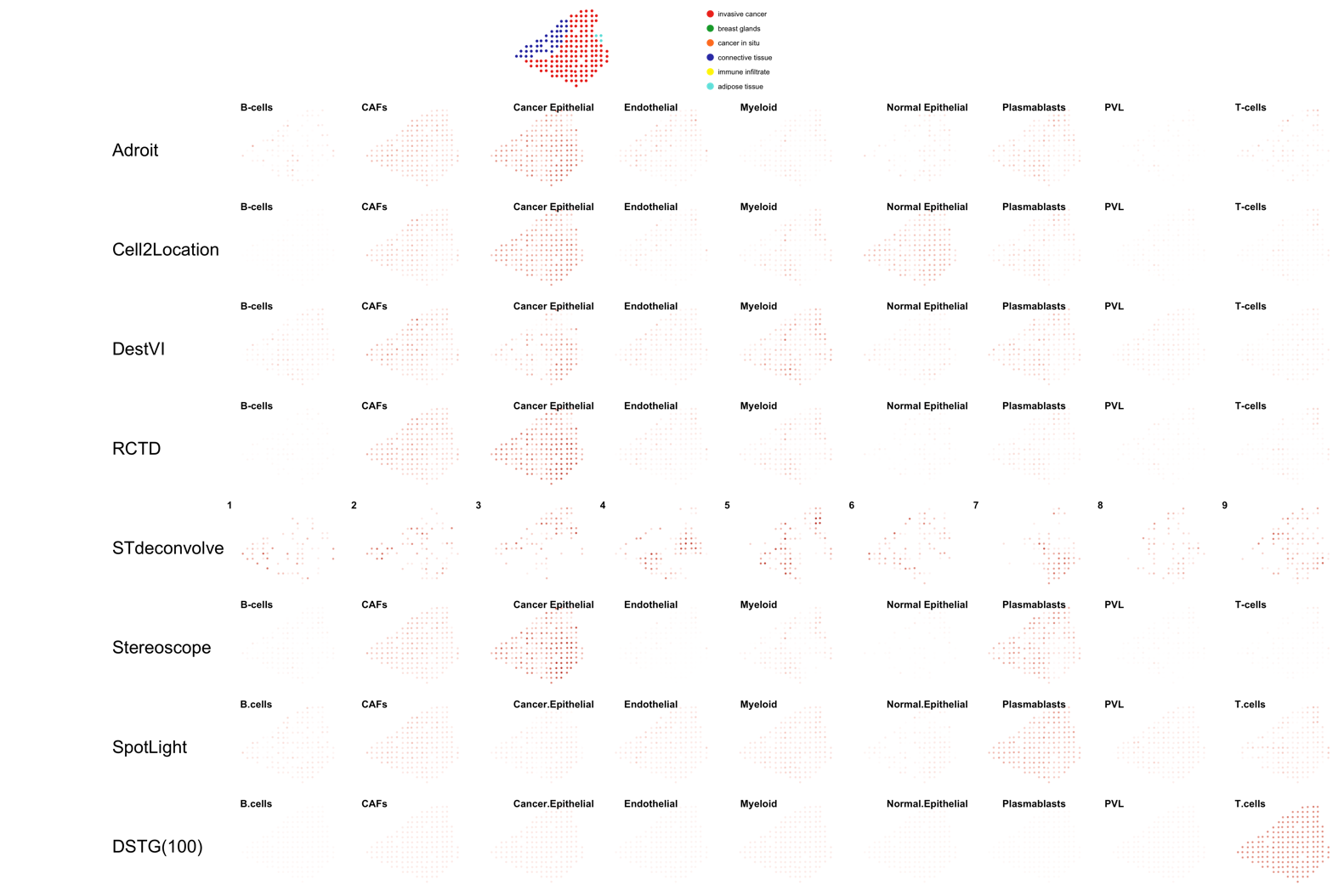

Supplementary Figure 17. Normalized cell type proportion estimates of the C1 slide. The first row is the pathologist annotated layer. Cell type proportion estimates are normalized, to sum up to 1. White indicates 0 and red indicates 1 such that the redder the color, the closer to a proportion of 1.

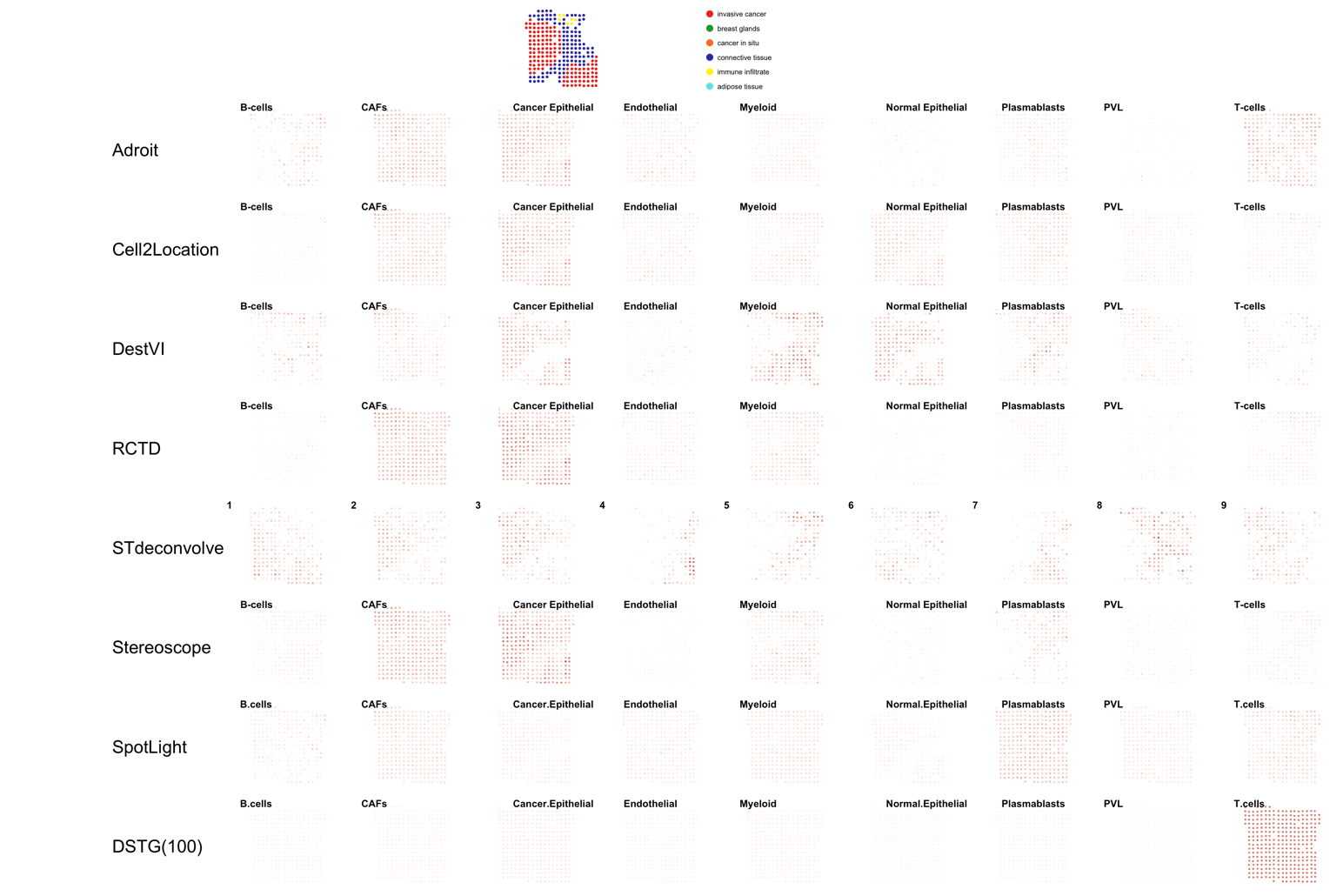

Supplementary Figure 18. Normalized cell type proportion estimates of the D1 slide. The first row is the pathologist annotated layer. Cell type proportion estimates are normalized, to sum up to 1. White indicates 0 and red indicates 1 such that the redder the color, the closer to a proportion of 1.

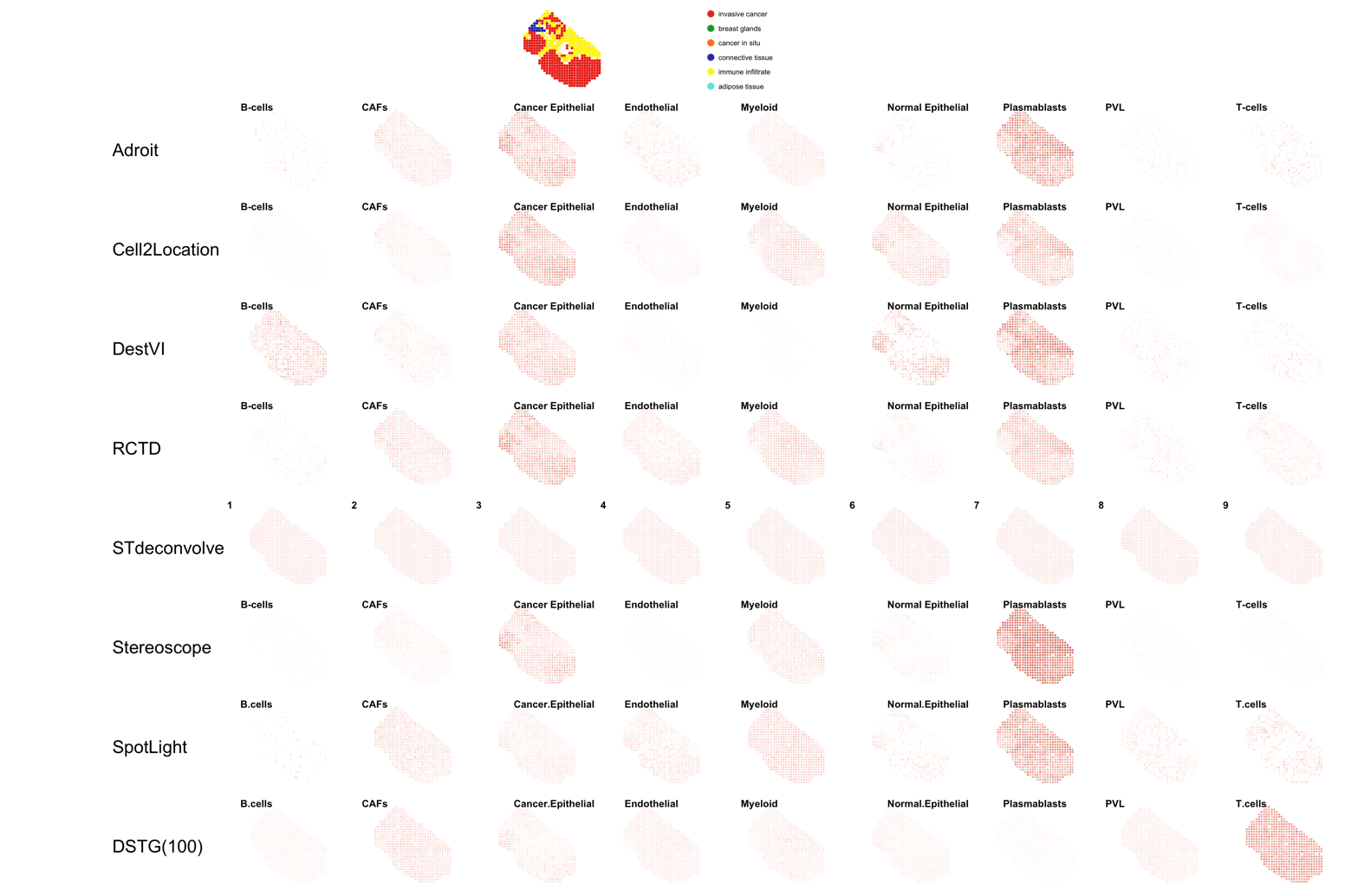

Supplementary Figure 19. Normalized cell type proportion estimates of the E1 slide. The first row is the pathologist annotated layer. Cell type proportion estimates are normalized, to sum up to 1. White indicates 0 and red indicates 1 such that the redder the color, the closer to a proportion of 1.

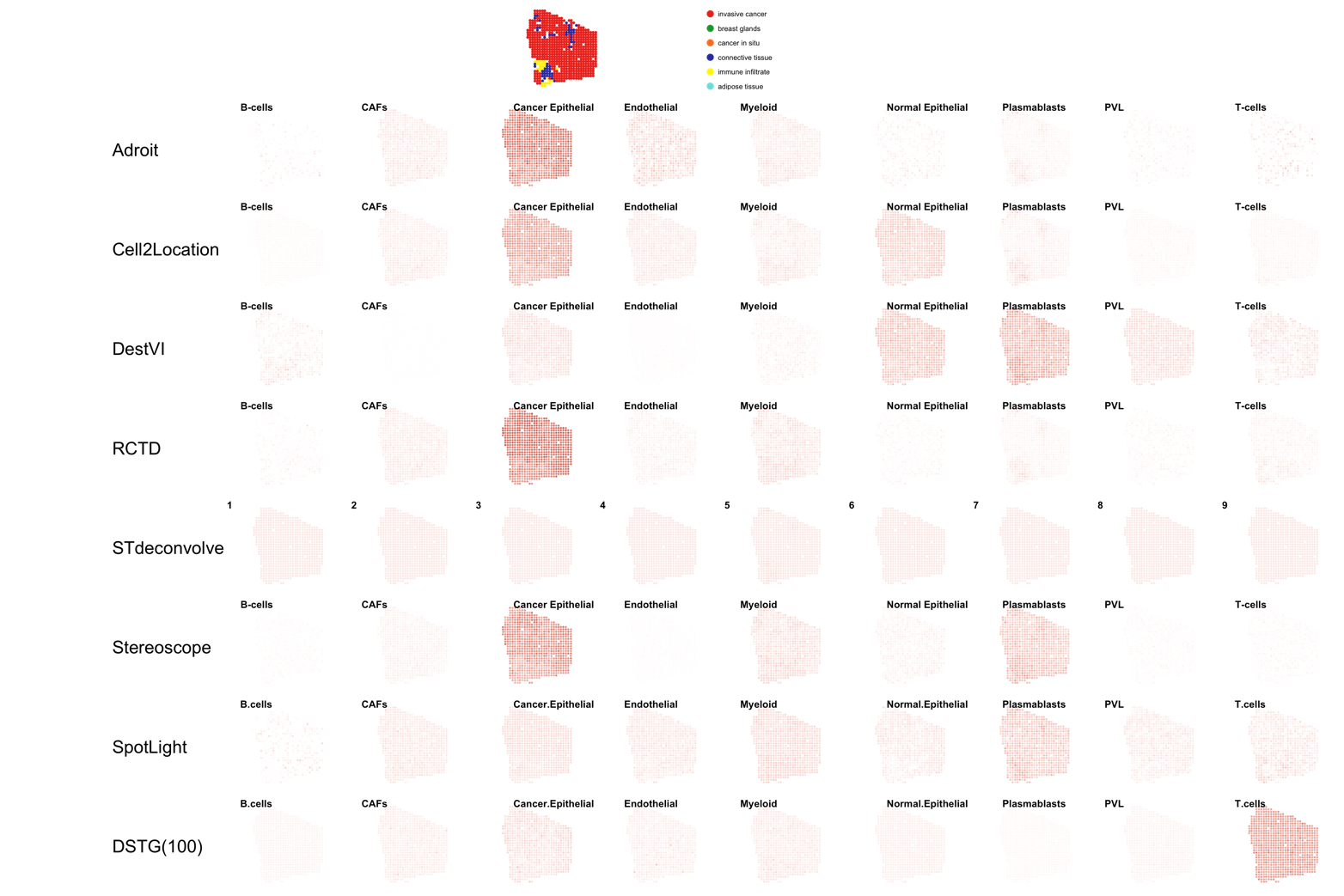

Supplementary Figure 20. Normalized cell type proportion estimates of the F1 slide. The first row is the pathologist annotated layer. Cell type proportion estimates are normalized, to sum up to 1. White indicates 0 and red indicates 1 such that the redder the color, the closer to a proportion of 1.

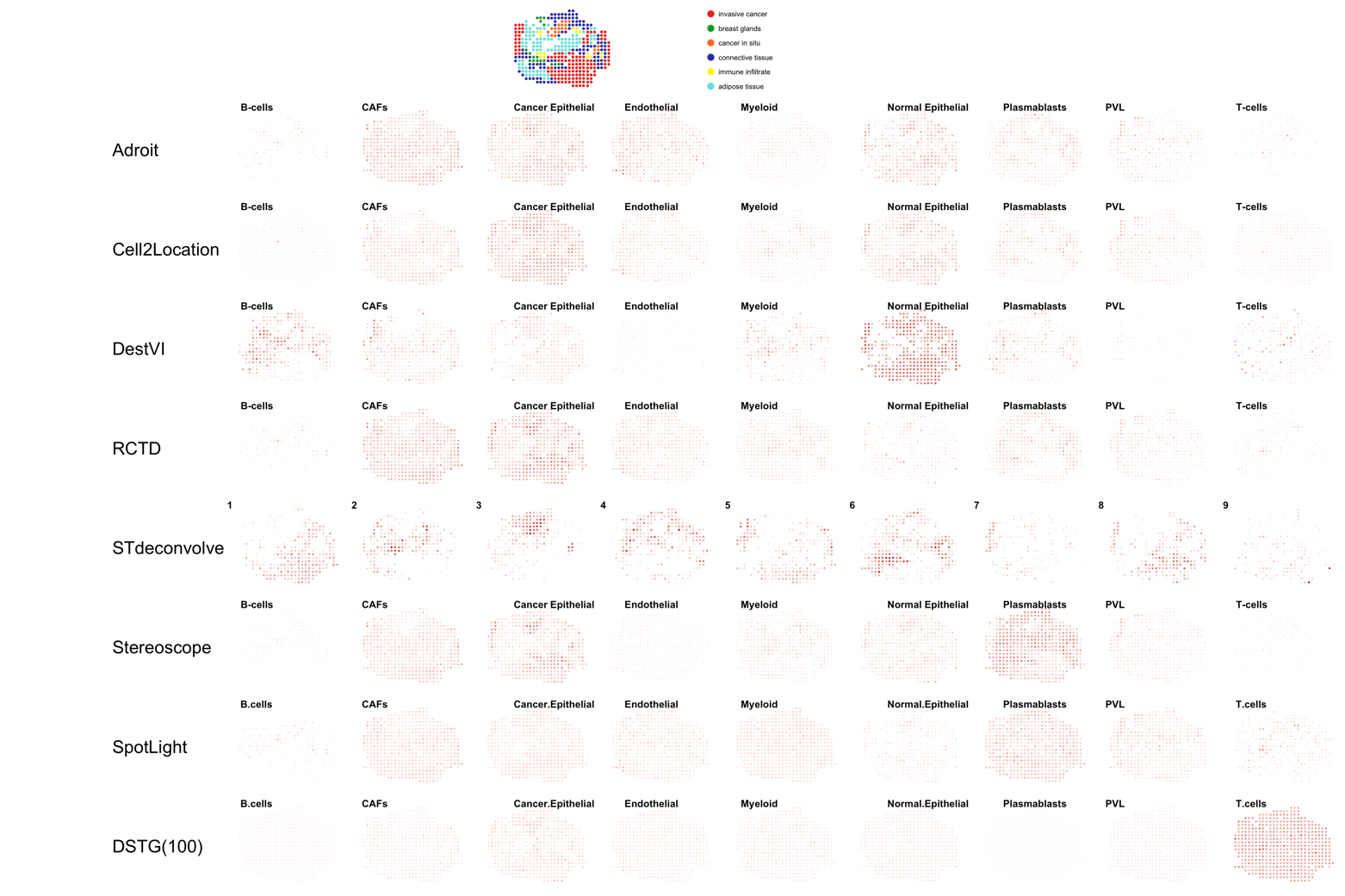

Supplementary Figure 21. Normalized cell type proportion estimates of the G2 slide. The first row is the pathologist annotated layer. Cell type proportion estimates are normalized, to sum up to 1. White indicates 0 and red indicates 1 such that the redder the color, the closer to a proportion of 1.

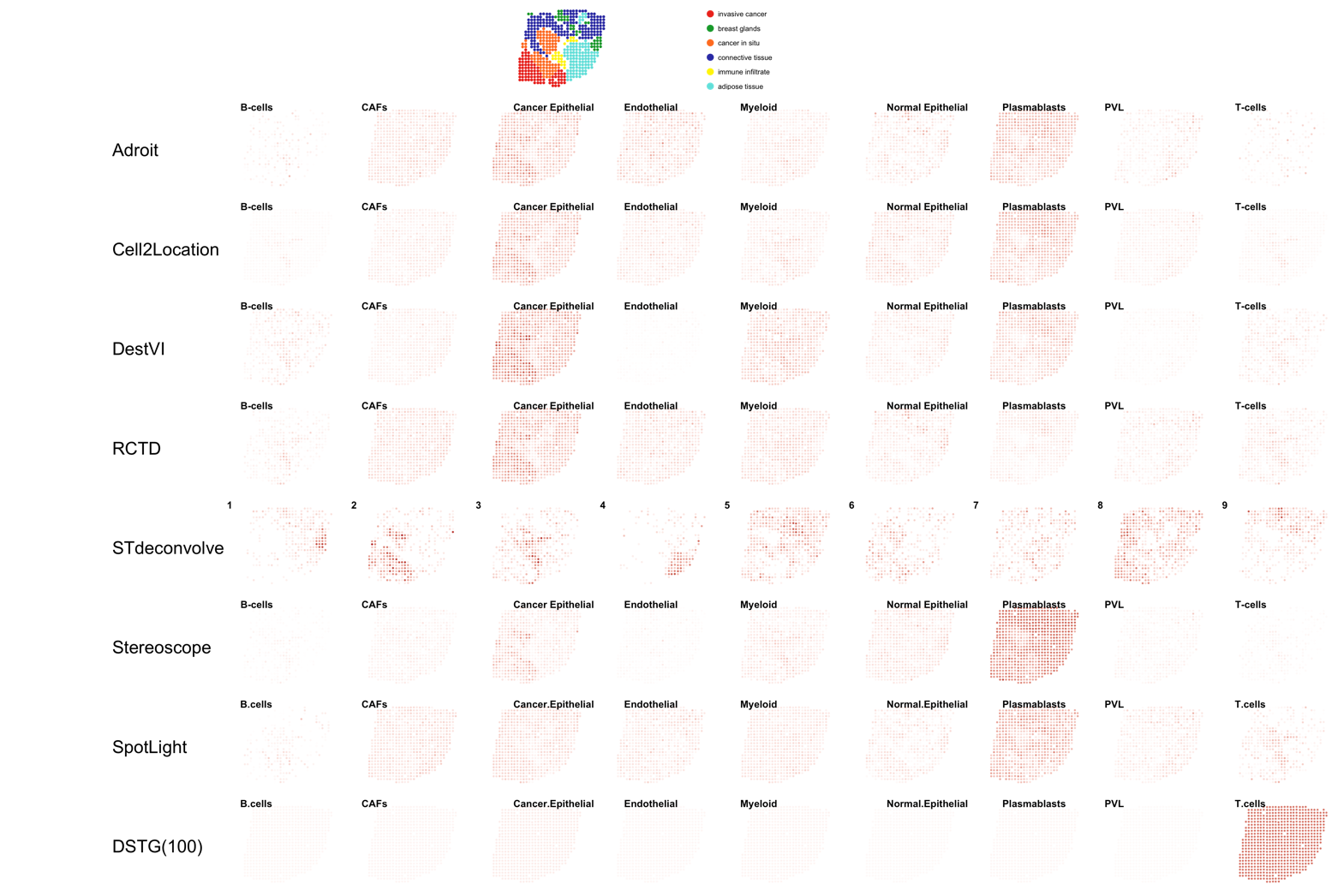

Supplementary Figure 22. Normalized cell type proportion estimates of the H1 slide. The first row is the pathologist annotated layer. Cell type proportion estimates are normalized, to sum up to 1. White indicates 0 and red indicates 1 such that the redder the color, the closer to a proportion of 1.
